# Proteasome subunit *PSMC3* variants cause neurosensory syndrome combining deafness and cataract due to proteotoxic stress

**DOI:** 10.1101/836015

**Authors:** Ariane Kröll-Hermi, Frédéric Ebstein, Corinne Stoetzel, Véronique Geoffroy, Elise Schaefer, Sophie Scheidecker, Séverine Bär, Masanari Takamiya, Koichi Kawakami, Barbara A. Zieba, Fouzia Studer, Valerie Pelletier, Claude Speeg-Schatz, Vincent Laugel, Dan Lipsker, Florian Sandron, Steven McGinn, Anne Boland, Jean-François Deleuze, Lauriane Kuhn, Johana Chicher, Philippe Hammann, Sylvie Friant, Christelle Etard, Elke Krüger, Jean Muller, Uwe Strähle, Hélène Dollfus

**Author notes:** equal contributors (Ariane Kröll-Hermi, Frédéric Ebstein, Corinne Stoetzel, Véronique Geoffroy).

## Abstract

Whole-genome sequencing was performed on patients with severe deafness and early-onset cataracts as part of a neurological, sensorial and cutaneous novel syndrome. A unique deep intronic homozygous variant in the *PSMC3* gene (c.1127+337A>G, p.Ser376Argfs15*), encoding the 26S proteasome ATPase ring subunit 5 (Rpt5) was identified leading to the transcription of a cryptic exon. Patient’s fibroblasts exhibited impaired protein homeostasis characterized by accumulation of ubiquitinated proteins suggesting severe proteotoxic stress. Indeed, the TCF11/Nrf1 transcriptional pathway allowing proteasome recovery after proteasome inhibition is permanently activated in the patient’s fibroblasts. Upon chemical proteasome inhibition this pathway is however impaired. These cells were unable to compensate for proteotoxic stress although a higher proteasome content. Two different zebrafish studies led to inner ear development anomalies as well as cataracts. PSMC3 proteasome subunit dysfunction leads to various neurological manifestations, early onset cataracts and deafness and suggest that Rpt5 plays a major role in inner ear, lens and central nervous system development.

## INTRODUCTION

Early onset deafness is one of the most common causes of developmental disorder in children (prevalence rate of 2-4/1000 infants) and identically early-onset cataract is the most important cause of paediatric visual impairment worldwide (prevalence form 2-13.6/10000 according to regions) accounting for 10% of the causes of childhood blindness. Each condition can be attributed to environmental causes (intrauterine infections, inflammation, trauma or metabolic diseases) or to genetic causes with a well-recognized very high level of genetic heterogeneity with 59 known genes causing early-onset cataracts and 196 genes known to cause severe deafness (Reis and Semina, 2018; Azaiez et al., 2018). Patients presenting both entities simultaneously, early onset severe deafness and congenital cataracts, are thought to be mainly due to teratogenic exposure during pregnancy especially infections and are, nowadays, considered to be very rare. Indeed, only very few genetic inherited entities associating both congenital cataracts and deafness have been reported so far. The Aymé-Gripp syndrome (cataract, deafness, intellectual disability, seizures and Down syndrome like facies) has been recently linked to *de novo* pathogenic variants in the *MAF* gene a leucine zipper-containing transcription factor of the AP1 superfamily. (Niceta et al., 2015) In addition, dominant pathogenic variants in *WFS1* (recessive loss of function variants are responsible for Wolfram syndrome) have been described in children with congenital cataracts and congenital deafness presenting in the context of neonatal/infancy -onset diabetes (De Franco et al., 2017).

Herein, using whole-genome sequencing, we describe a novel homozygous non-coding pathogenic variant in *PSMC3* associated with severe congenital deafness and early onset cataracts and various neurological features in 3 patients from a very large consanguineous family. *PSMC3* encodes the 26S regulatory subunit 6A also known as the 26S proteasome AAA-ATPase subunit (Rpt5) of the 19S proteasome complex responsible for recognition, unfolding and translocation of substrates into the 20S proteolytic cavity of the proteasome(TANAKA, 2009). The proteasome is a multiprotein complex involved in the ATP-dependent degradation of ubiquitinated proteins to maintain cellular protein homeostasis and to control the abundance of many regulatory molecules. In mammalian cells, a major compensation mechanism for proteasome dysfunction is governed by the ER membrane-resident TCF11/Nrf1 protein (Steffen et al., 2010; Radhakrishnan et al., 2010; Sotzny et al., 2016). Typical stimuli for TCF11/Nrf1 activation include proteasome inhibition and/or impairment which results in the release of C-terminal processed TCF11/Nrf1 fragment from the ER membrane following a complex series of molecular events involving the enzymes NGLY1 and DDI2. The cleaved TCF11/Nrf1 fragment enters then into the nucleus and acts as a transcription factor to promote the expression of ARE-responsive genes including 19S and 20S proteasome subunits, thereby augmenting the pool of proteasomes so that protein homeostasis can be preserved. (Steffen et al., 2010; Radhakrishnan et al., 2010; Sotzny et al., 2016) We suggest that biallelic loss of PSMC3 causes a novel autosomal recessive syndrome with varying degrees of neurosensorial dysfunctions including the combination of cataract and deafness. Functional analysis of patient’s cells revealed that although normal amount of proteasome proteins can be observed in steady-state conditions, the cells are unable to adapt to proteotoxic stress. The use of zebrafish morpholinos and CRIPSR-Cas9 assays confirmed the same combination of sensory phenotypes upon inactivating PSMC3 expression.

## RESULTS

### Patient phenotypes

Three patients with a novel syndromic neuro-sensory-cutaneous presentation consulted independently to our clinical centre over a period of 15 years. Careful analysis revealed that they originated from the same small village (Amarat) in the Kayseri region of Turkey and belong to the same large extended consanguineous family (Figure 1A). The proband is a male individual (II.4) diagnosed at the age of 7 months with severe perceptive deafness and subsequently benefited from a cochlear implantation. He was referred at the age of 2 years old to our centre because of visual impairment due to bilateral cataracts for which he underwent bilateral lensectomies. With years, he developed severe developmental delay and severe intellectual deficiency (no words, limited comprehension). Several facial features were noted (Table 1 and Figure 1B). In addition, autistic features and peripheral polyneuropathy of lower limbs were revealed at the age of 2.5 years old (Table 1). A full metabolic exploration was normal. At the age of 5 he developed sub cutaneous deposits at the level of the knees and elbows (Figure 1C). At the age of ten he developed white hair at the level of the two legs as opposed to the dark pigmented hair on the rest of the body. The 2 other patients (II.2 and II.7) were referred at the age of one year old and share the same severe congenital perceptive deafness (for which they also benefited from a cochlear implantation), visual impairment due to bilateral obstruent cataracts (for which they also had bilateral lensectomies) and sub cutaneous deposits. Patient II.2 did not present with autistic features but moderate developmental delay (read and write few words, no comprehension of complex sentences). Patient II.7 did not have peripheral polyneuropathy of lower limbs compared to the other patients.

**Table 1.**
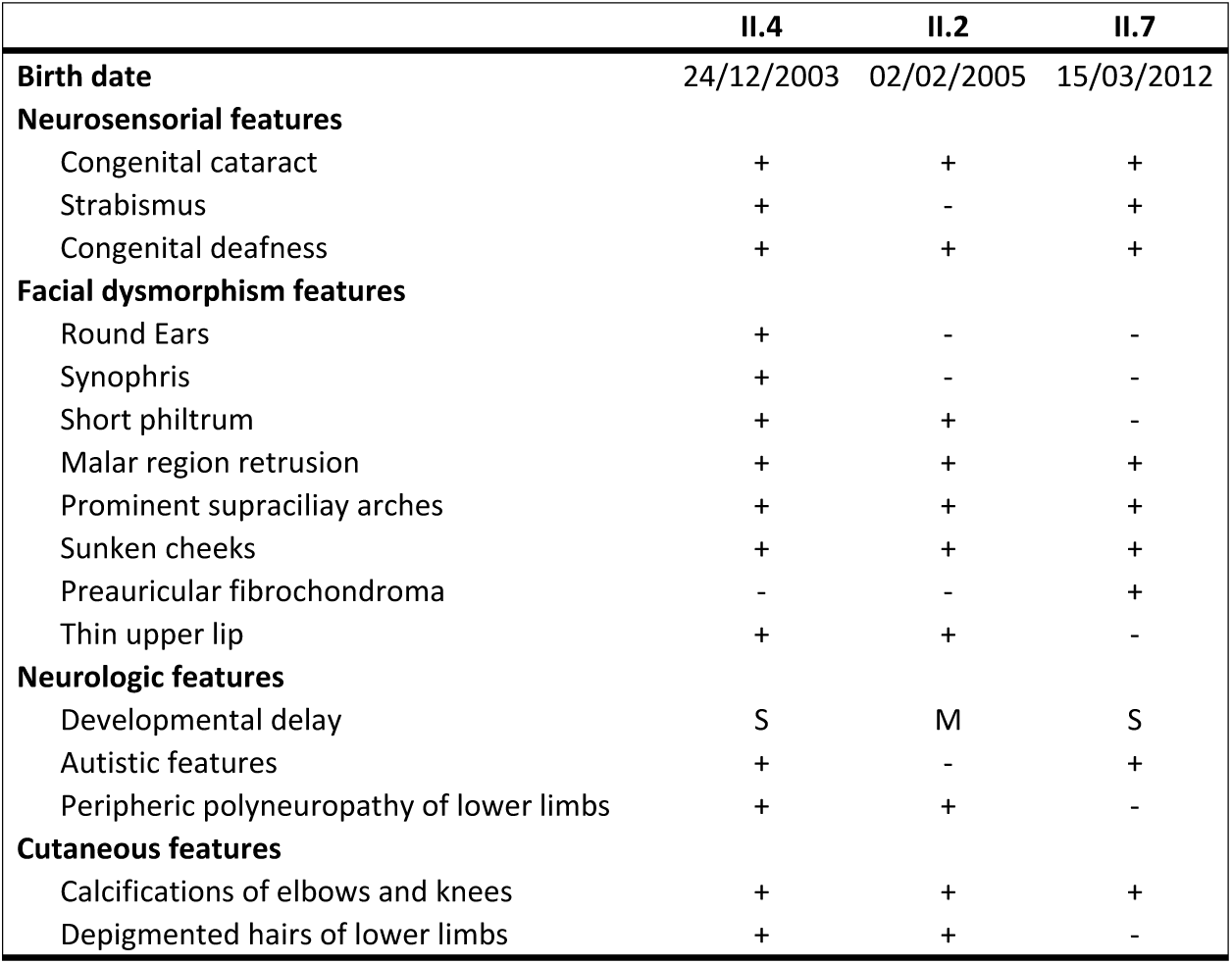
Clinical description of the patients with *PSMC3* pathogenic variants. M: moderate, S: severe

**Figure 1.**
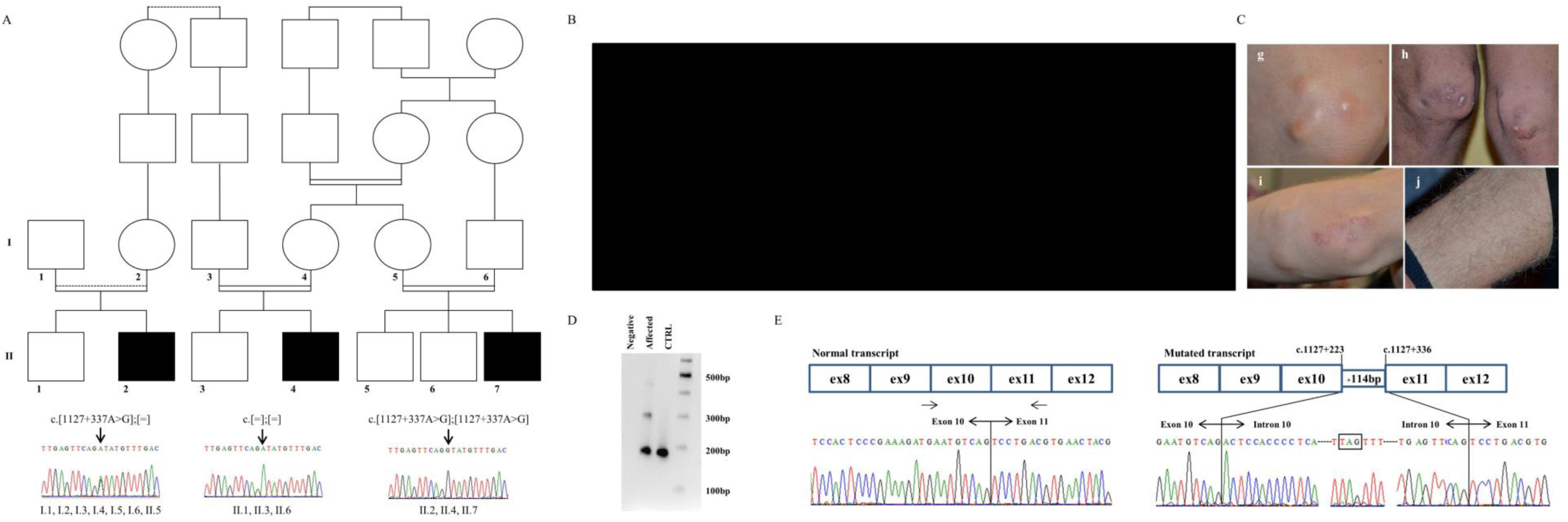
Family pedigree and cDNA analysis. (**A**) Family pedigree. Variant segregation analysis of *PSMC3*. Electropherogram of a part of intron 10 of *PSMC3* encompassing the identified variation (c.[1127+337A>G];[1127+337A>G], p.[(Ser376Arg15*)];[(Ser376Arg15*)]) in the affected individuals, their unaffected parents and siblings. The variation was found at the homozygous state in the affected individuals (II.2, II.4, II.7), at the heterozygous state in the parents (I.1, I.2, I.3, I.4, I.5, I.6) and was either at the heterozygous state (II.5) or absent in the unaffected siblings (II.1, II.3, II.6). (**B**) Face (up) and profile (down) photographs for patients II.4 (a: 8 yo, b: 16 yo), II.2 (c: 6 yo, d: 14 yo) and II.7 (e: 1 yo, f: 7 yo) over time. One can observe prominent supraciliay arches, synophris, sunken cheeks, short philtrum and retrusion in the malar region. (**C**) Sub cutaneous calcifications found only on knees (g: 9 yo and h: 16 yo) on elbows (i: 9 yo) of patient II.4. White hair were present only on the legs of the 3 patients as illustrated for patient II.4 (j: 16 yo). (**D**) Amplification of the cDNA fragment between exons 9-10 and 11 of *PSMC3* showing the abnormally spliced RNA fragment. One band at 180bp representing the normal allele is seen for the control and two bands for the individual II.4 (pathologic allele at 300bp). (**E**) Schematic representation for the incorporation of the 114bp intronic sequence resulting from the c.1127+337A>G deep intronic variation on the mRNA. Sanger sequencing of the fragment between exons 9-10 and 11 of the *PSMC3* cDNA obtained from patient II.4 fibroblasts’ RNA, showing the insertion of the 114bp cryptic exon. As a comparison, the schematic representation and sequence from a control individual are shown below.

### Identification of a rare deep intronic variation in *PSMC3*

During the years of follow-up each patient was explored for known deafness and cataract genes by Sanger sequencing (in particular for *GJB2*, one of the patient being an heterozygous carrier of the c.30delG well known pathogenic variant) but also using larger assays such as Whole-Exome Sequencing (WES) with a specific focus on known deafness and cataract genes (Table S1) and standard chromosomal explorations (karyotype and chromosomal microarray analysis) but all were negative (see supplementary methods). Considering that affected individuals may harbour pathogenic variants in a region not covered by the WES (i.e. intronic, intergenic…) or not well detected (i.e. structural variations)(Geoffroy et al., 2018b), we applied whole-genome sequencing (WGS) to the three affected individuals (II.2, II.4 and II.7) and two healthy individuals (II.1 and II.3). Given the known consanguinity in the family, our analysis was focused on homozygous variations and more specifically within homozygous regions defined by the SNP arrays (Figure S1). In addition to the classical filtering strategy including functional criteria, frequency in population based databases and co-segregation analysis (see methods), we defined a list of 4846 potentially interacting genes with the already known human cataract (59) and deafness genes (196) (Figure S2). This strategy allowed us to identify from the ∼5,000,000 variations per WGS, 6 variations out of which a unique homozygous variant in the intron 10 of the *PSMC3* gene (c.1127+337A>G, p.?) remained of interest (Table S2 and Figure S3). This variant was not present in any variation database (e.g. gnomAD) and is predicted to create a new donor splice site. Interestingly among others PSMC3 was shown to interact (Table S3 and S4, Figure S4) with CHMP4B (MIM 610897), ACTG1 (MIM 102560) and GJB6 (MIM 604418) involved in cataract and deafness.

### Effect of the variation on *PSMC3* expression and localisation

In order to assess the effect of this deep intronic variation, we investigated the expression of the gene in the patient’s fibroblasts. The suspected new donor site could be associated to multiple acceptor sites within intron 10 (Figure S5). RNA analysis revealed an additional band specific to the affected patient that was further explored by Sanger sequencing (Figure 1D). The consequence of this variation is the inclusion of a cryptic 114 bp exon during the splicing process based on the intronic sequence (r.1127_1127+1insACTCCACCCCTCATCTGAAGGCACAGAGGCTGGAGGCACTTAGTTTCCTGGCCTCACACC TCAGCCCATTAACACACGCCAGGAATGGCCGGGACCAGATGGACTTGAGTTCAG) (Figure 1E) that is predicted to add 15aa (LeuHisProSerSerGluGlyThrGluAlaGlyGlyThr) at position 376 followed by a stop (p.(Ser376Argfs15*)). Analysis at the RNA level showed a significantly reduced level of *PSMC3* mRNA as well as the presence of an additional truncated form (Figure S6). However, no difference in expression or localisation of the PSMC3 protein could be detected between the control and the patient cells under normal condition indicating that the truncated form is probably not stable (Figure S7).

### Functional effect of the PSMC3 variant to the proteasome function and assembly

Given the role of PSMC3 in protein degradation, we determined the intracellular level of ubiquitinated proteins in patient cells compared to controls. Our results show an increased level of ubiquitinated proteins in patient cells (Figure 2A and B), suggesting that the proteasomal proteolysis is less efficient. Having shown a possible effect on proteasome function, we next investigated how the variant could affect the proteasome assembly and dynamics. First, in standard condition, PSMC3 protein and its partners were immunoprecipitated from either controls’ or patient’s fibroblasts and revealed by mass spectrometry (Figure 2C). The PSMC3 protein was detected in the input control and patient indicating that the variant does not affect the protein stability of the remaining wild type allele confirming the western-blot analysis. Looking at the interacting partners, one can notice that each proteasome subunit could be detected revealing no apparent defect in the general organisation of the proteasome. However, protein abundance of each proteasome subcomplex estimated from the number of mass spectrometry spectra observed between the controls and the patient (Table S5) shows a general increase of the proteasome subunits (approximatively 20%). Looking more specifically at each sub complex, the increase is mainly due to the core particle including the alpha and beta rings with 1.5 fold for all PSMA proteins and most PSMB protein and even a 2.0 fold increase for PSMB2/4/6 while PSMC and PSMD remain at the same ratio.

**Figure 2.**
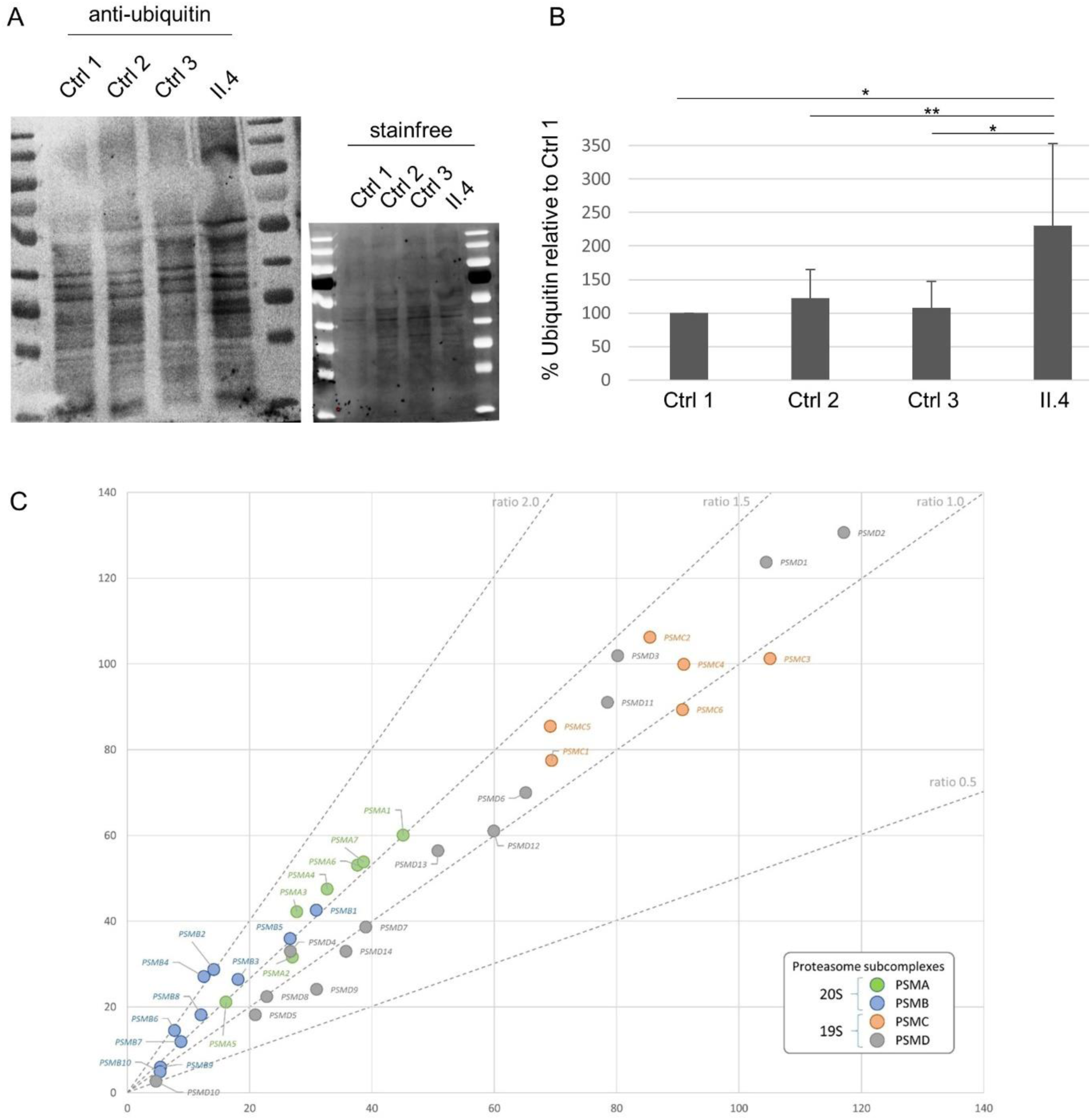
Effect of the deep intronic PSCM3 variant on the proteasome function. (**A**) Anti-ubiquitin western blot in control and patient fibroblasts and total amount of proteins loaded (stainfree) showing increased ubiquitination in patient cells (lane 4). (**B**) Histogram showing the quantification of ubiquitin with western blot assays. The data shown correspond to the sum of all bands detected by the anti-ubiquitin antibody expressed as a percentage of the amount of ubiquitin in “Control 1” cells. Results are the mean of ten independent experiments. *: p<0.01, **: p<0.05. (**C**) Mass spectrometry results from the co immunoprecipitation with PSMC3 are displayed as the normalized total number of spectra count of each protein computed as the mean from 3 controls (x axis) vs the mean of patient II.4 triplicate. Proteasome subcomplexes are colored according to the displayed legend and standard ratio lines are drawn.

To further characterize the consequences of the *PSMC3* variant on the functionality of the ubiquitin-proteasome system (UPS), cell lysates derived from control and patient primary fibroblasts were analysed by non-denaturing PAGE with proteasome bands being visualized by their ability to hydrolyse the fluorogenic peptide. As shown in Figure 3A, gel overlay assay for peptidase activity revealed two strongly stained bands corresponding to the positions of the 20S and 26S proteasome complexes, respectively. However, no discernible differences could be detected in the chymotrypsin-like activity of both 20S and 26S complexes between control and patient cells, suggesting that peptide hydrolysis in the 20S proteolytic core is not substantially impaired by the *PSMC3* pathogenic variant. The notion that proteasome activity does not vary between these two samples was further confirmed by monitoring the degradation rate of the Suc-LLVY peptide directly in whole-cell extracts from control and patient fibroblasts over a 180-min period that was almost identical in both samples (Figure 3B). In order to characterize the proteasome populations in cells carrying the deep intronic *PSMC3* homozygous variant, samples separated by non-denaturing PAGE were subsequently analysed by western blotting. As expected, using an antibody against the proteasome subunit Alpha6, two major bands were observed in the 20S and 26S regions (Figure 3C). Interestingly, the signal intensity for both proteasome complexes was significantly stronger in patient fibroblasts, suggesting that the amount of intracellular proteasome pools in these cells was higher than those of control fibroblasts. Western blotting against the PSMC3 subunit revealed two bands in the 26S proteasome area and corresponding to single and double-capped proteasomes (19S-20S and 19S-20S-19S, respectively) and confirmed the higher amount of these complexes in patient cells (Figure 3C), however there are some lower bands corresponding to 19S precursor intermediates indicating that assembly of 19S complexes is affected. As shown in Figure 3C, staining for PA28-α, a subunit of the alternative proteasome regulator PA28, revealed one major band corresponding to the position of the 20S proteasome, indicating the 20S proteasomes in these samples mainly consist of PA28-20S complexes. Again, patient fibroblasts exhibited a stronger expression level of such homo-PA28 complexes than their wild-type counterparts. To validate the notion that the homozygous pathogenic variant results in increased assembly of newly synthetized proteasome complexes, patient fibroblasts were compared to control ones for their content in various proteasome subunits using SDS-PAGE followed by western-blotting. As illustrated in Figure 3D, the steady-state expression level of most of the β, α and Rpt subunits was substantially higher in the patient’s cells. Unexpectedly, the increased proteasome content was accompanied by a parallel rise of ubiquitin-modified proteins in these cells, suggesting that proteasomes from patients bearing the homozygous *PSMC3* pathogenic variant, although being in greater number, are ineffective. Altogether, these results point to a defective proteasome function in subjects carrying the deep intronic homozygous *PSMC3* pathogenic variant, which seems to be compensated by an ongoing assembly of newly synthetized 20S and 26S complexes.

**Figure 3.**
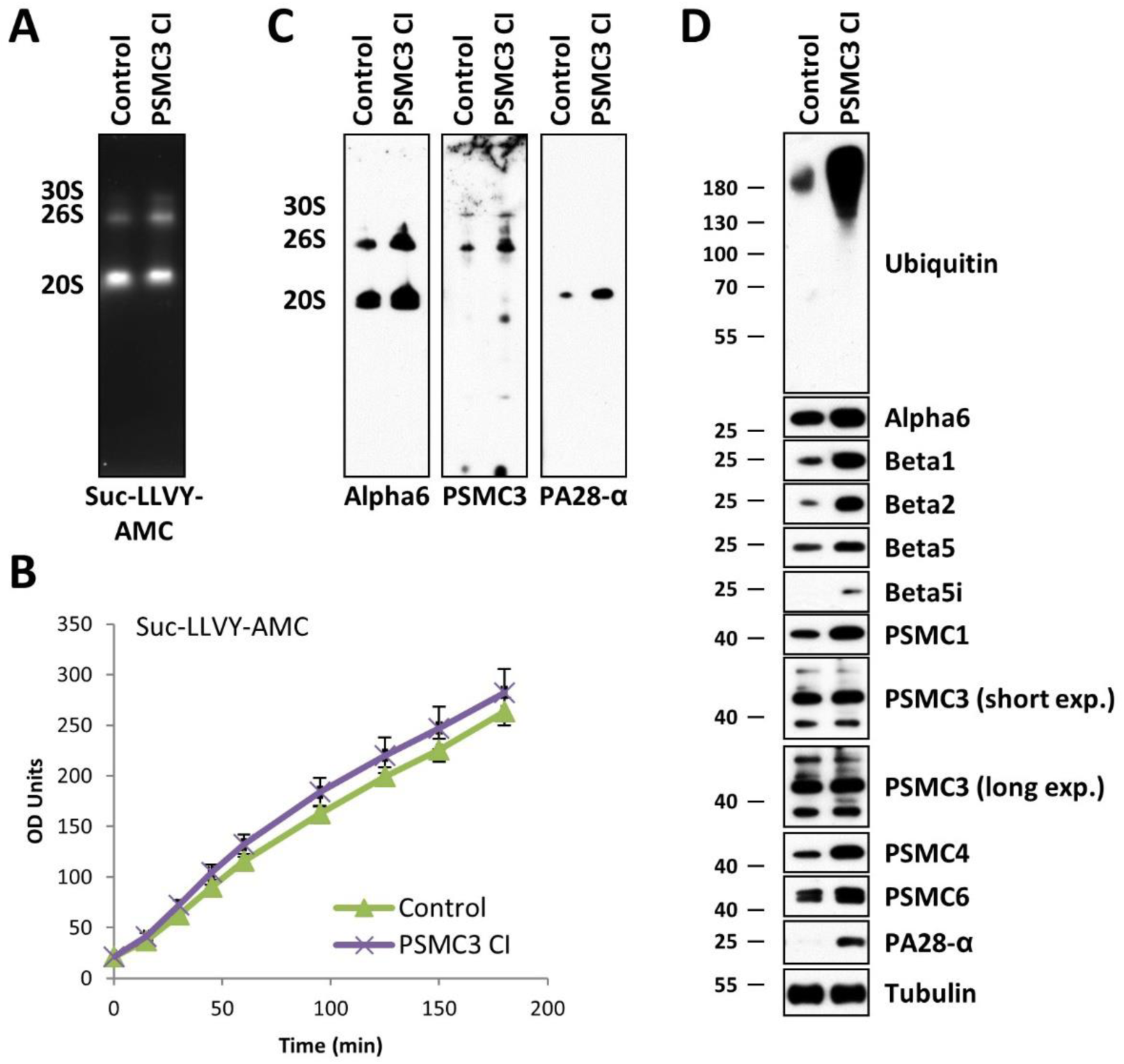
Fibroblasts derived from patient carrying the c.1127+337A>G homozygous *PSMC3* variation exhibit an increased amount of both proteasome complexes and ubiquitin-protein conjugates. (**A**) Whole-cell lysates from control and patient (case index, CI) fibroblasts were assessed by 3-12% native-PAGE gradient gels with proteasome bands (30S, 26S and 20S complexes) visualized by their ability to cleave the Suc-LLVY-AMC fluorogenic peptide. (**B**) Ten micrograms of control and patient cell lysates were tested for their chymotrypsin-like activity by incubating them with 0.1 mM of the Suc-LLVY-AMC substrate at 37°C over a 180-h period of time in quadruplicates on a 96-well plate. Indicated on the y-axis are the raw fluorescence values measured by a microplate reader and reflecting the AMC cleavage from the peptide. (**C**) Proteasome complexes from control and patient fibroblasts separated by native-PAGE were subjected to western-blotting using antibodies specific for Alpha6, Rpt5 (PSMC3) and PA28-α, as indicated. (**D**) Proteins extracted from control and CI PSMC3 were separated by 10 or 12.5% SDS-PAGE prior to western-blotting using primary antibodies directed against ubiquitin and several proteasome subunits and/or components including α6, β1, β; β5, β5i, Rpt2 (PSMC1), Rpt5 (PSMC3), Rpt3 (PSMC4), Rpt4 (PSMC6) and PA28-α, as indicated. For the PSMC3 staining, two exposure times are shown. Equal protein loading between samples was ensured by probing the membrane with an anti-α-Tubulin antibody.

We next sought to determine the impact of this variant on the ability of the cells to respond to perturbed protein homeostasis following proteasome dysfunction. To this end, both control and patient cells were subjected to a 16-h treatment with the β5/β5i-specific inhibitor carfilzomib in a non-toxic concentration prior to SDS-PAGE and western-blotting analysis using various antibodies specific for proteasome subunits. As shown in Figure 4, control cells exposed to carfilzomib could successfully compensate the applied proteotoxic stress by increasing their pools of intracellular proteasomes, as evidenced by elevated expression of all investigated β- and Rpt-subunits. As expected, this process was preceded by the processing of the ER membrane-resident protein TCF11/Nrf1 (Figure 4A), which is the transcription factor acting on nuclear genes encoding 19S and 20S proteasome subunits (Steffen et al., 2010; Radhakrishnan et al., 2010; Sotzny et al., 2016). Strikingly, the level of processed TCF11/Nrf1 in response to carfilzomib was much lower in cells carrying the homozygous *PSMC3* pathogenic variant than that observed in control cells. Accordingly, the patient’s fibroblasts were unable to upregulate their proteasome subunits following proteasome inhibition, as determined by decreased expression levels of the βsubunits (Figure 4A). Altogether these data indicate that, although exhibiting higher amounts of proteasomes under normal conditions, fibroblasts derived from patients carrying the homozygous *PSMC3* pathogenic variant fail to preserve protein homeostasis under stress conditions, as a result of proteasome deficiency.

**Figure 4.**
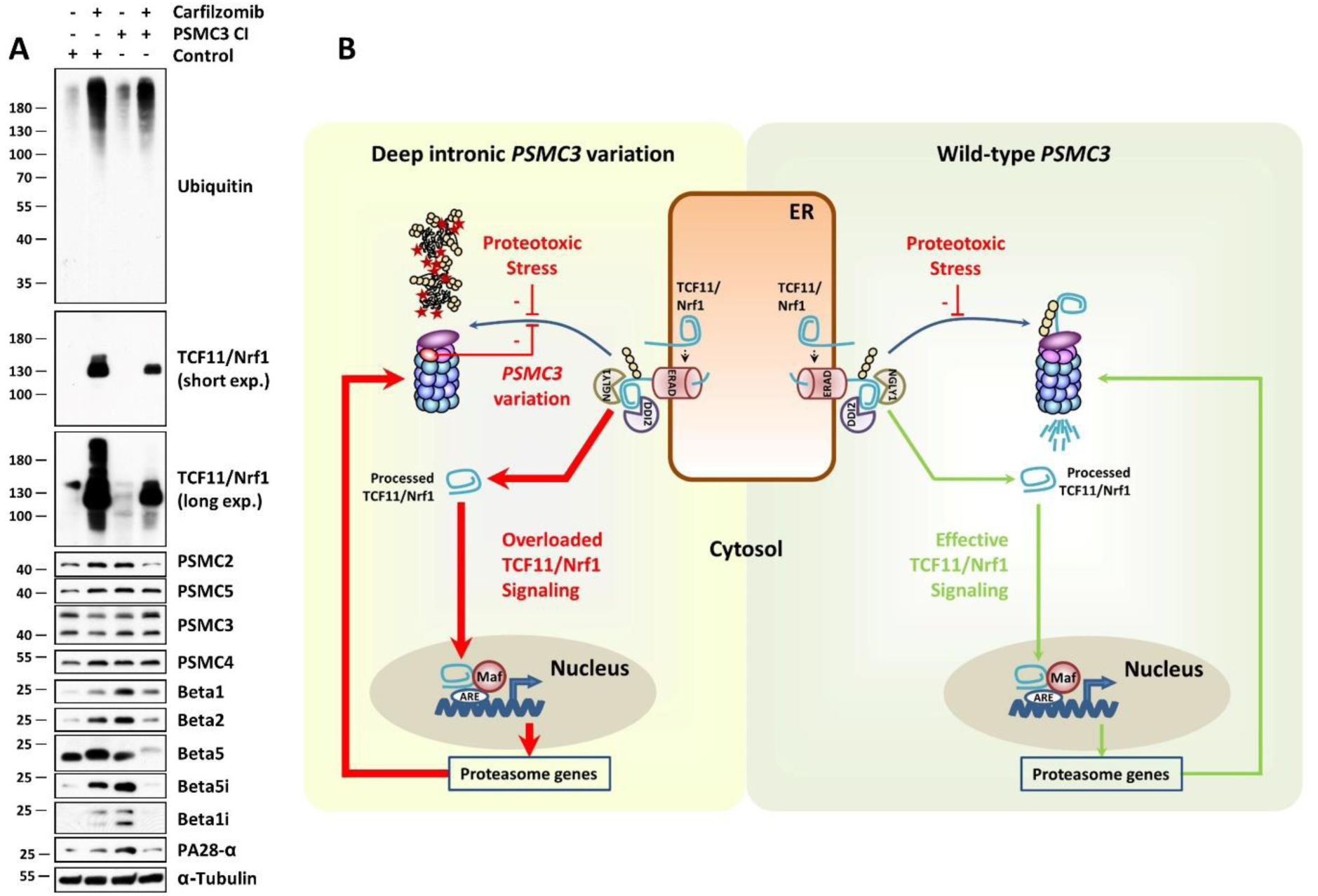
Patient fibroblasts carrying the c.1127+337A>G homozygous *PSMC3* variation exhibit an exhausted TCF11/Nrf1 processing pathway which prevents them to upregulate proteasome subunits in response to proteotoxic stress. **(A)** Control and patient (index case, CI PSMC3) fibroblasts were exposed to a 16-h treatment with 30 nM of the proteasome inhibitor carfilzomib or left untreated (as a negative control). Following treatment, cells were collected and subjected to RIPA-mediated protein extraction prior to SDS-PAGE and subsequent western-blotting using antibodies specific for ubiquitin, TCF11/Nrf1, Rpt1 (PSMC2), Rpt3 (PSMC4), Rpt5 (PSMC3), Rpt6 (PSMC5), β1, β2, β5, β5i, β1i, PA28-α and α-Tubulin (loading control) as indicated. For the TCF11/Nrf1 staining, two exposure times are shown. (**B**) Schematic diagram depicting TCF11/Nrf1 processing pathway in response to proteasome dysfunction and/or proteotoxic stress in patient carrying the deep intronic homozygous *PSMC3* variation (left) and in heathy subjects (right). Under normal conditions, TCF11/Nrf1 is a short-lived ER-membrane (endoplasmic reticulum) resident protein, which is rapidly subjected to proteasome-mediated degradation following retro-translocation to the cytosol by ER-associated degradation machinery (ERAD). In case of proteasome dysfunction (i.e. proteotoxic stress), the half-life of TCF11/Nrf1 is prolonged and become then a substrate for the NGLY1 and DDI2, thereby giving rise to a C-terminal cleaved fragment that enters into the nucleus. After nuclear translocation, cleaved TCF11/Nrf1 associates with Maf and promotes the expression of proteasomes genes, so that protein homeostasis can be restored. In patients carrying the c.1127+337A>G homozygous *PSMC3* variation, defective proteasomes promote a constitutive activation of TCF11/Nrf1 thereby resulting in increased assembly of newly synthetized non-functional proteasomes, which in turn activate TCF11/Nrf1 again. Over-activation of TCF11/Nrf1 results in pathway exhaustion thus rendering patient cells incapable of responding to further proteotoxic stress.

### Effect of PSMC3 loss of function in zebrafish similar to patients’ phenotype

To establish a functional link between the observed decreased proteasome activity and the phenotype observed in the patients, we next assessed the lens and the ear in the zebrafish model. The zebrafish orthologue (Ensembl, ENSDARG00000007141) of human *PSMC3* is located on chromosome 7 with two predicted protein-coding splice variants (404 and 427 amino acids). Both zebrafish *psmc3* isoforms share 83% sequence identity with the human orthologue. We confirmed by *in situ* hybridization that *psmc3* is maternally expressed and not spatially restricted (Figure S8) (Thisse and Thisse, 2004). Injection of morpholinos against *psmc3* generated zebrafish morphants embryos that were examined at 4 days post-fertilization (dpf) for lens or ear abnormalities. The lens size of morphants was slightly smaller than in control or uninjected embryos (Figure S9A). For cataract detection, we used a protocol based on confocal reflection microscopy, a labelling-free non-invasive imaging method that enables the detection of abnormal light reflection in the lens of living embryos (Figure 5A).(Takamiya et al., 2016) Morpholino injections revealed significant abnormal lens reflections in 95% of the morphants (n=55), whereas only 2% of 5 bp mismatched morpholino injected controls (n=45) and none of the uninjected embryos (n=20) showed cataract (Figure 5B-B’).

**Figure 5.**
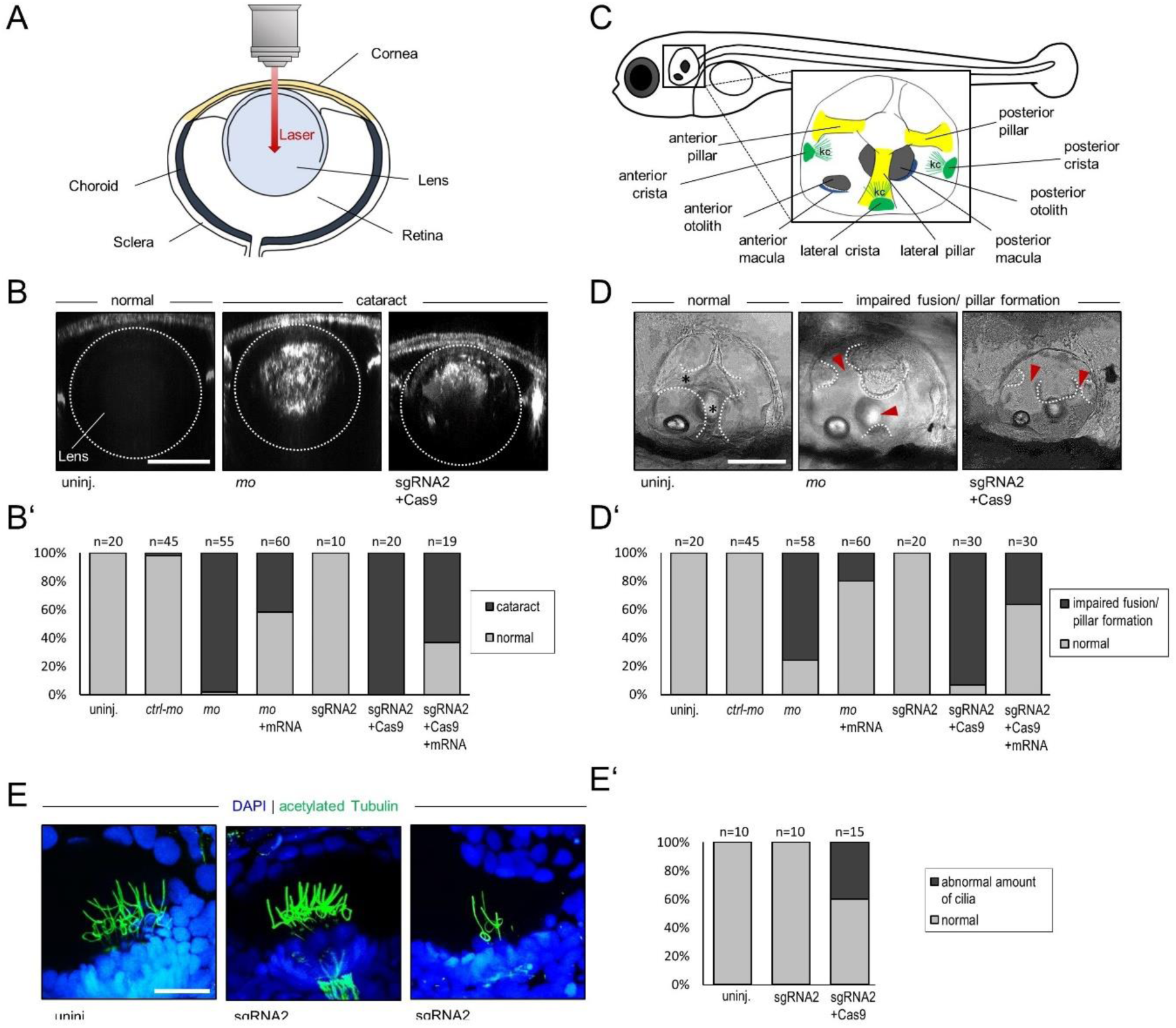
*psmc3* morphants and F0 mosaic zebrafish exhibit cataract and show abnormalities during the semicircular canal development in the ear. **(A)** Scheme of a zebrafish eye. (**B-B’**). Cataract detection revealed abnormal lens reflection in *psmc3* morpholino (MO)-mediated knockdown but not in controls (uninj, *ctrl-mo*). Similarly, abnormal lens reflection was also observed in embryos injected with sgRNA + Cas9 but not in sgRNA injected embryos without Cas9 (sgRNA2). Co-injection of wt *psmc3* mRNA with either *psmc3-mo* or sgRNA2 + Cas9 reduced the number of embryos presenting abnormal lens reflection. Scale bar = 50 µm. (**B’**) Quantification of embryos with abnormal lens reflection. (**C**) Representative image of a zebrafish ear at 4 dpf. kc= kinocilia. (**D-D’**) bright field images of inner ear development (lateral position). (**D**) Epithelial projections were fused and formed canal pillars in 4-day-old uninjected and control injected fish (*ctrl-mo*, sgRNA2) but not in morphants (*mo*) and crispants (sgRNA2+Cas9). Co-injection of wt *psmc3* mRNA with *psmc3-mo* or sgRNA + Cas9 reduced the number of embryos presenting abnormal ear phenotype. Black asterisks indicate fused pillars. Red arrowheads mark unfused projections. Scale bar = 100 µm. (**D’**) Quantification of embryos with abnormal projection outgrowth. (**E**) An anti-acetylated tubulin antibody (green) staining revealed an abnormal amount of kinocilia in *psmc3* crispants (sgRNA2+Cas9) compared to uninjected and control injected embryos (sgRNA2). Nuclei are stained in blue with DAPI. Representative images show kinocilia of the lateral cristae. Scale bar = 20 µm. (**E’**) Quantification of embryos with an abnormal amount of kinocilia.

As deafness has been reported for all three human patients, we subsequently examined the inner ear development of 4 dpf zebrafish morphants (Figure 5C). *psmc3* morphants displayed a smaller ear compared to control or uninjected embryos (Figure S9C). Interestingly, the majority of morphants presented anomalies during the semicircular canal morphogenesis. While the epithelium projections of all uninjected (n=20) and all control-injected (n=45) embryos were fused and formed pillars after 4 dpf, the canal projections of morphants failed to fuse in 79% of the cases (n=58, Figure 5D-D’). The specificity of the morpholino was confirmed by a rescue experiment by co-injecting the full-length splice morpholino resistant zebrafish *psmc3* mRNA. The cataract phenotype was rescued in ∼58% of the cases (n=60) while the ear phenotype was rescued in 76% of the cases (n=60) (Figure 5B’+D’). The outgrowth of the epithelial projections of the developing anterior and posterior semicircular canals begins around 48 hpf. From 57-68 hpf, the projections fuse in the center of the ear and form the pillars. This is followed by the outgrowth of the projection of the lateral semicircular canal around 57 hpf and completed by the fusion with the other two pillars in the center around 70 hpf.(Geng et al., 2013) To analyze the semicircular canal morphogenesis in *psmc3* morphants, life imaging was performed between 56 and 72 hpf in the transgenic line gSAIzGFF539A expressing a GFP signal in all three pillars. In all morphants (n=9), the projections failed to fuse and form pillars during the observed time frame. In two morphants (22%), the outgrowth of epithelial projections even failed completely. In contrast, the projections of uninjected (n=2) and control injected (n=2) embryos fused during the observed period in the center of the ear and formed canal pillars (Movie S1). Previous studies showed that a reduced number or a smaller size of otoliths, crystal-like structures required for the transmission of mechanical stimuli to the hair cells, can lead to deafness and balancing difficulties in zebrafish.(Han et al., 2011; Stooke-Vaughan et al., 2015) *psmc3* morphants did not present any otolith defect (Figure S10A-C). In addition, expression of *otopetrin*, a gene required for proper otolith formation, was unaffected at 28 hpf and 4 dpf (Figure S9D).

As autism has been reported for one of the patients and since brain malformation have been reported previously in some autistic patients and autistic zebrafish morphants (Elsen et al., 2009), we investigated possible morphological brain changes in *psmc3* morphants. *In-situ* hybridization targeting brain markers such as *krox20, msxc, her8a* and *sox19b* was performed on 24 hpf embryos but did not reveal any obvious differences in their expression patterns (Figure S11).

To confirm these results, we additionally used the CRISPR/Cas9 system to knockdown *psmc3* (Figure 5B-D). The high cutting efficiency of the CRISPR/Cas9 founders (F0) were evaluated to 55.2% (gRNA1) and 53.7% (gRNA2) (Figure S12).(Etard et al., 2017) CRISPR/Cas9 founders (i.e. crispants) are often genetically mosaics. However, in cases of highly efficient gRNAs, they have been shown to recapitulate mutant phenotypes successfully.(Teboul et al., 2017; Küry et al., 2017; Paone et al., 2018) Both *psmc3* crispants used in this study displayed a cataract phenotype (100% of gRNA1+Cas9 (n=20) and 95% of gRNA2+Cas9 (n=20), whereas none of the control embryos (injected with gRNA1 (n=20) or gRNA2 (n=10) without the Cas9 protein) showed abnormal lens reflections (Figure 5B-B’ and S13A-A’). Moreover, we recapitulated the ear phenotype seen in *psmc3* morphants. While the ear pillars of uninjected (n=10) and control-injected (n=20) embryos formed after 4 dpf, the projections of both crispants failed to fuse in 65% of gRNA1 (n=10) and 93% of gRNA2 (n= 30) injected embryos (Figure 5D-D’ and S12B-B’). Co-injection of gRNA2, Cas9 and the *psmc3* mRNA also led to a partial rescue of the lens and ear phenotype with only 63% of abnormal lens reflection instead of 100% (n=19) and 37% of unfused canal projections instead of 93% (n=30) (Figure 5D’). Performing an *in situ* hybridization examining the mRNA expression of *versican a* and *versican b*, two genes suggested to be required for a proper canal fusion event (Geng et al., 2013), significant differences could be observed after 72 hpf. Indeed, both genes were highly expressed in the whole ear tissue, whereas *versican a* was not expressed and *versican b* was restricted to the dorsolateral septum in wildtype or control injected embryos (Figure S13B-D).

As kinocilia, specialized microtubule-based structures found on the surface of hair cells, have been shown to play a key role at least in mechanosensation during development(Kindt et al., 2012), we immunostained crispants (sgRNA2 injected with Cas9) and control injected embryos (sgRNA2 without Cas9) at 5 dpf using anti-acetylated tubulin antibody. In 40% of crispants (n=15), a reduced number of kinocilia was observed while the number of control injected embryos (n=10) was similar to that of uninjected embryos (n=10) (Figure 5E+E’).

Taken together, these zebrafish assays confirm that *psmc3* plays a very important role in the development of a transparent lens and the semicircular canals of the inner ear - reminiscent of the human phenotype described herein.

## DISCUSSION

In this study, we describe a novel homozygous deep intronic splice variant, identified in 3 patients with an unusual neurosensorial disease combining early onset deafness, cataracts and sub cutaneous deposits, in *PSMC3* encoding one of the proteasome subunit. Clinical data and functional analysis in patient’s cells and zebrafish proved the effect of the variation and the consequences leading to a recessive form of proteasome deficit with a haploinsufficiency mechanism. The observation of a neurosensory disease broadens the spectrum of ubiquitin-proteasome system (UPS) related disorders. Sequencing the entire genome of patients gives access to the whole spectrum of their variations and possibly disease causing ones. WGS is a powerful tool (Belkadi et al., 2015) helping to identify variations not covered or missed by WES such as structural variations(Geoffroy et al., 2018b) or deep intronic variations (Vaz-Drago et al., 2017). Interestingly, in this study, we combined to WGS, homozygosity mapping and *in silico* predicted interactors to narrow down to the region of *PSMC3*. The 3 patients carried an homozygous deep intronic variation (i.e. >100 bases from the exon-introns boundaries)(Vaz-Drago et al., 2017) with a predicted splicing effect on the *PSMC3* gene that we confirmed on the patient’s cells. We then focused on demonstrating the effect of this variation at the level of the proteasome.

The 26S proteasome, the chief site of protein turnover in eukaryotic cells, consists of 2 complexes: a catalytic 28-subunit barrel shaped core particle (20S) that is capped at the top or the bottom by one 19 subunit regulatory particle (19S). The core particle contains the catalytic subunits β1, β2 and β5 exhibiting caspase-, trypsin- and chymotrypsin-like activities, respectively. Recognition of a substrate with the requisite number and configuration of ubiquitin is mediated principally by both Rpn10 and Rpn13 subunits which act as ubiquitin receptors (Husnjak et al., 2008; Deveraux et al., 1994). To allow substrate degradation, ubiquitin is first removed by Rpn11, a metalloprotease subunit in the lid (Yao and Cohen, 2002). The globular domains of a substrate are then unfolded mechanically by a ring-like heterohexameric complex consisting of six distinct subunits, Rpt1 to Rpt6, which belong to the ATPases-associated-with-diverse-cellular-activities (AAA) family.(Chen et al., 2016) PSMC3 encodes for Rpt5 involved in the substrate unfolding and translocation, which are then presumably catalysed. (TANAKA, 2009; Lam et al., 2002)

To our knowledge, this is the first report of a human biallelic pathogenic variant occurring in one of the ATPase Rpt subunits of the base of the 19S regulatory particle. Recently, *de novo* pathogenic variations in the non-ATPase subunit *PSMD12* (Rpn5) of the 19S regulator lid of the 26S complex have been reported in 6 patients with neurodevelopmental disorders including mainly intellectual disability (ID), congenital malformations, ophthalmic anomalies (no cataracts), feeding difficulties, deafness (unspecified type for 2 patients/6), and subtle facial features.(Küry et al., 2017) *PSMD12* variants have been also associated to a large family with ID and autism and 1 simplex case with periventricular nodular heterotopia.(Khalil et al., 2018) *PSMD12* is highly intolerant to loss of function (LoF) variations and the most likely effect is haploinsufficiency due to the *de novo* heterozygote occurrence of loss of function truncating, non-sense or deletion variants. Interestingly, according to the gnomAD and DDD data(Karczewski et al., 2019; Huang et al., 2010), *PSMC3* is also predicted to be extremely intolerant to LoF variation. Indeed, the haploinsufficiency score of *PSMC3* is 4.76 that is within the high ranked genes (e.g. HI ranges from 0% to 10%) from the DECIPHER data. The pLI score (0.96) makes it among the highest intolerant genes (e.g. a score >0.9 defines the highest range) with only 3 observed LoF variants versus 23.2 predicted and confidence interval = 0.13). This explains also maybe the rarity of LoF variations found to date in this gene. In our cases, this is the first time that biallelic class 5 variations are reported in one of the proteasome subunit delineating a recessive mode of inheritance. Nevertheless, the fact that the homozygous variation is affecting the splicing machinery and leads to a reduced but not abolished expression of *PSMC3* could mimic a possible haploinsufficiency mechanism. Both proteomic and biochemical approaches undertaken in this study revealed that the deep intronic homozygous *PSMC3* variation is associated with increased amounts of 26S and 20S-PA28 proteasome complexes (Figures 2C and 3). The observation that patient cells concomitantly increase their intracellular pool of ubiquitin-protein conjugates (Figures. 2A and 3D) is surprising and strongly suggests that such proteasomes are defective. In support of this notion, we found that, although carrying greater amounts of proteasomes, patient fibroblasts did not exhibit higher chymotrypsin-like activity compared to control cells, which can be at least partly explained by upregulation of PA28 and the concomitant increased peptide-hydrolysis (Ma et al., 1992) (Figures 3A and B). The C-terminus of Rpt5, which is supposed to be truncated in the patient due to the deep intronic splice variant, has been shown to be important for gate opening of the α-ring of the 20S proteasome core complex and for assembly of the 19S complex (Smith et al., 2007; Singh et al., 2014). Thus, an expression of this truncated Rpt5 variant even in low mounts may disturb proteasome assembly and function. These data led us to conclude that the increased steady-state expression level of the proteasome subunits observed in patient fibroblasts might reflect a constitutive *de novo* synthesis of proteasomes, which aims to compensate the dysfunctional ones. Strikingly and in contrast to control cells, TCF11/Nrf1 is constitutively processed in patient cells (Figure 4A), confirming that patients’ proteasomes were impaired. This, in turn, gives rise to a pathological vicious circle of events in which TCF11/Nrf1 and defective proteasomes stimulate each other (Figure 4B). We reasoned that such a process may result in a pathway overload which in turn reduce the ability of TCF11/Nrf1 to respond to further proteotoxic stress. Consistent with this hypothesis and unlike control cells, patient fibroblasts were not capable of upregulating their proteasome subunits when challenged with proteasome inhibitor carfilzomib (Figure 4A). This result is of great importance, as it demonstrates that, under stress conditions, cells carrying the deep intronic homozygous *PSMC3* variation suffer from proteasome deficiency. Because cataract and semicircular canal malformations are observed in zebrafish embryos depleted with *PSMC3* -and *a fortiori* proteasomes-, these data established a clear cause-and-effect relationship between the deep intronic *PSMC3* variant and the acquisition of patients’ phenotype. On the other hand, one cannot exclude that the pathogenesis of the homozygous *PSMC3* variation may involve additional mechanisms. Because target genes of TCF11/Nrf1 include anti-inflammatory factors (Yang et al., 2018; Widenmaier et al., 2017), it is also conceivable that inflammation might play a role in this process. This assumption would be in line with the observation that subjects suffering from other loss-of-function proteasome variations such as *PSMB8* (Agarwal et al., 2010; Liu et al., 2012; Arima et al., 2011), *PSMA3, PSMB4, PSMB9* (Brehm et al., 2015) and/or *POMP* (Poli et al., 2018) exhibit an inflammatory phenotype including joint contractures. In any case, the potential contribution of innate immunity to the pathogenesis of the homozygous *PSMC3* variant via TCF11/Nrf1 warrants further investigation (Agarwal et al., 2010; Arima et al., 2011).

The ubiquitin-proteasome system (UPS) is a protein degradation pathway that regulates the intracellular level of proteins involved in a very wide variety of eukaryote cellular functions. Thus, it is not surprising that this pathway is related to multiple human conditions including cataract, where an overburden of malfunctional and aggregated proteins cannot be adequately removed by the UPS (Shang and Taylor, 2012). Moreover, protein degradation dysfunction is recognized as a widespread cause of neurodegenerative diseases such as Parkinson, Huntington or Alzheimer diseases. Several inherited rare disorders have been shown to be related to directly UPS dysfunction in enzymes of the ubiquitin conjugation machinery: *UBE3A* in Angelman syndrome(Kishino et al., 1997, 3), *UBE2A* in X-linked syndromic ID(Nascimento et al., 2006), *UBE3B* in Kaufmann oculo cerebrofacial syndrome (classified as blepharophimosis – mental retardation syndrome) or other related enzymes such as *HUWE1* an ubiquitin ligase in X-linked for dominant ID syndrome(Froyen et al., 2008) (Moortgat et al., 2018) and *OTUD6B* a deubiquitinating enzyme for neurodevelopmental disability with seizures and dysmorphic features(Santiago-Sim et al., 2017). The phenotypic tropism for neurodevelopmental disorders especially with intellectual disability and other brain dysfunctions (autism, seizures, brain malformations) seem to be overall a hallmark of the gene alterations related with the UPS inherited dysfunctions. In the family presented herein, all 3 patients had various neurodevelopmental anomalies (autism, ataxia, mild ID) and mild facial dysmorphism.

However, the striking clinical presentation of the cases reported herein is the combination of early onset deafness and early onset cataracts that motivated (independently) the referral of the 3 cases. These features have not yet been related to this group of disorders to date. Moreover this association has only been reported twice in larger syndromic forms such as the dominant form of *WFS1* but in the context of neonatal/infancy -onset diabetes (De Franco et al., 2017) and the Aymé-Gripp syndrome (including also intellectual disability, seizures and Down syndrome like facies) with *de novo* pathogenic variants in the *MAF* gene (Niceta et al., 2015). Variations in the later one were shown to impair *in vitro* MAF phosphorylation, ubiquitination and proteasomal degradation.

The zebrafish model was extremely useful to demonstrate the effect of a reduced *psmc3* expression leading to inner ear anomalies and lens opacities. The overall development of the lens and inner ear as well as the underlying gene regulatory networks are highly conserved throughout evolution.(Cvekl and Zhang, 2017) Pathogenic variations or knockdown of genes implicated in human cataract and/or deafness in zebrafish have been shown to recapitulate similar phenotypes.(Gao et al., 2017; Mishra et al., 2018; Yousaf et al., 2018; Takamiya et al., 2016; Busch-Nentwich, 2004) Using two strategies (morpholino and CRISPR/Cas9) the fish developed a cataract and ear phenotype that could be rescued. We did not observe other obvious anomalies. Interestingly, a reduction of proteasome activity has previously been associated with lens defects in zebrafish. The knock out of the zebrafish gene *psmd6* and the knock down of *psmd6* and *psmc2*, both encoding for proteins of the UPS, resulted in a severe impairment of lens fibre development. The ear phenotype of the *psmd6* mutant and both morphants was not assessed.(Imai et al., 2010) A direct link between the UPS and auditory hair cell death or impaired semicircular canal morphogenesis has not been described in zebrafish yet. However, knock down of *atoh1*, a gene regulated by the UPS, has been shown to severely affect hair cell development in the inner ear of zebrafish.(Millimaki et al., 2007) The of canal pillars observed in zebrafish *psmc3* morphants and crispants might be also a secondary effect, as abnormal sensory cristae with few hair cells have been previously assumed to lead to an abnormal development of semicircular canals.(Cruz et al., 2009; Haddon and Lewis, 1991)

In conclusion, our work demonstrates the implication of a deep intronic variant in a novel ultra-rare neurosensorial syndrome with early onset cataract and deafness in one of the proteasome subunit, *PSMC3.* Although, *de novo* dominant variations have been associated to several proteasome related disorders, we report for the first time a bi allelic pathogenic variant. Our observations strongly suggest that the amount of PSMC3 is critically implied in the development and maintenance of the inner ear and the lens.

## MATERIALS AND METHODS

### Patients and ethics

The 3 cases have consulted independently and were enrolled subsequently by the CARGO (reference center for rare eye diseases at the Strasbourg University Hospital, France). All participants were assessed by a clinical geneticist, a neuro pediatrician, an ENT specialist, a dermatologist and a paediatric ophthalmologist. Written consent for research and publication was obtained for all study participants. This research followed the tenets of the Declaration of Helsinki. Approval was obtained from our institutional review board “*Comité Protection des Personnes*” (EST IV, N°DC-20142222). The identified gene was submitted to the GeneMatcher tool but no other patient with the same phenotype could be identified (Sobreira et al., 2015).

### Homozygosity regions

Three affected individuals (II.2, II.4, and II.7) and 2 unaffected individuals (II.1 and II.3) were analyzed with the Affymetrix GeneChip® Mapping 250K Array Xba 240 (Affymetrix, Santa Clara, CA). Sample processing and labelling were performed according to the manufacturer’s instructions. Arrays were hybridized on a GeneChip Hybridization Oven 640, washed with the GeneChip Fluidics Station 450 and scanned with a GeneChip Scanner 3000. Data were processed by the GeneChip DNA Analysis Software version 3.0.2 (GDAS) to generate SNP allele calls. An average call rate >99% was obtained. Homozygosity regions were identified as regions of homozygosity longer than 25 adjacent SNPs. Shared regions of homozygosity are visualized by the HomoSNP software (IGBMC, Strasbourg), which displays one patient per line (Figure S1). Three regions of homozygosity of respectively 0.3, 2.1 and 1.3 Mb on chromosome 11 (11:44,396,024-44,668,374; 11:45,574,574-47,684,908; 11:66,066,993-67,349,899) are shared between the affected and not the healthy individuals.

### Whole Exome Sequencing analysis

Whole exome sequencing (WES) was performed in 2014 for the three affected siblings (II.2, II.4, II.7) and one healthy brother (II.6) by the IGBMC (Institut de Génétique et de Biologie Moléculaire et Cellulaire, Illkirch-Graffenstaden, France) Microarray and Sequencing platform. Exons of DNA samples were captured using the in-solution enrichment methodology (Agilent SureSelect All Exon XT2 50 Mb Kit) and sequenced with an Illumina HiSeq 2500 instrument to generate 100 bp paired-end reads. The reads were mapped to the reference human genome (GRCh37/hg19) using the Burrows-Wheeler Aligner (BWA v0.7.1) (Li and Durbin, 2009). The HaplotypeCaller module of the Genome Analysis ToolKit (GATK v3.4-46) was used for calling both SNV and indel (DePristo et al., 2011). Structural Variations (SV) were called using CANOES (Backenroth et al., 2014).

From 96,222 to 102,072 genetic variants (SNV/indel/SV) were identified per proband from the WES analysis (Supplementary Table 1). Bioinformatic analyses (see Materials and methods) highlighted a unique homozygous missense variation in the *DGKZ* gene (NM_001105540.1:c.1834G>A, p.Ala612Thr) encoding for the Diacylglycerol Kinase Zeta, located in a homozygous region of interest on chromosome 11 (Table S1). This variant, reported previously with a gnomAD frequency of 0.018%, was predicted tolerated by SIFT(Ng and Henikoff, 2003) and neutral by PolyPhen-2 (Adzhubei et al., 2010) and was finally manually ruled out, as we were unable to explain the patients phenotype based on the gene function.

### Whole Genome Sequencing (WGS)

Whole Genome Sequencing was performed in 2016 for the three affected siblings (II.2, II.4, II.7) and two healthy brothers (II.1, II.6) by the *Centre National de Recherche sur le Génome Humain* (CNRGH, Evry France). Genomic DNA was used to prepare a library for WGS using the Illumina TruSeq DNA PCR-Free Library Preparation Kit. After normalization and quality control, qualified libraries were sequenced on a HiSeq2000 platform (Illumina Inc., CA, USA), as paired-end 100 bp reads. At least 3 lanes of HiSeq2000 flow cell were produced for each sample, in order to reach an average sequencing depth of 30x. Sequence quality parameters were assessed throughout the sequencing run and standard bioinformatics analysis of sequencing data was based on the Illumina pipeline to generate FASTQ files for each sample. The sequence reads were aligned to the reference sequence of the human genome (GRCh37) using the Burrows-Wheeler Aligner (BWA V7.12) (Li and Durbin, 2009). The UnifiedGenotyper and HaplotypeCaller modules of the Genome Analysis ToolKit (GATK) (DePristo et al., 2011), Platypus (https://github.com/andyrimmer/Platypus) and Samtools (Li et al., 2009) were used for calling both SNV and indel. Structural Variations (SV) were called using SoftSV (Bartenhagen and Dugas, 2016). Moreover, each known cataract and deafness genes were visually inspected with IGV (Thorvaldsdóttir et al., 2013).

### Bioinformatics analysis

Annotation and ranking of SNVs/indels and structural variations were performed respectively by VaRank (Geoffroy et al., 2015) (in combination with Alamut Batch, Interactive Biosoftware, Rouen, France) and by AnnotSV (Geoffroy et al., 2018a). Variant effect on the nearest splice site was predicted using MaxEntScan (Yeo and Burge, 2004), NNSplice (Reese et al., 1997) and Splice Site Finder (Shapiro and Senapathy, 1987). Very stringent filtering criteria were applied to filter out non-pathogenic variants: (i) variants represented with an allele frequency of more than 1% in public variation databases including the 1000 Genomes (1000 Genomes Project Consortium et al., 2015), the gnomAD database (Lek et al., 2016), the DGV (MacDonald et al., 2014) or our internal exome database, (ii) variants in 5′ and 3′ UTR, downstream, upstream, intronic and synonymous locations without pathogenic prediction of local splice effect. The *PSMC3* nomenclature is based on the accession number NM_002804.4 from the RefSeq database (O’Leary et al., 2016). Genomic coordinates are defined according to GRCh37/hg19 assembly downloaded from the University of California Santa Cruz (UCSC) genome browser. (Tyner et al., 2017).

### Sanger confirmation and segregation

The variant confirmation and the cosegregation analysis with the phenotype in the family member were performed by Sanger sequencing after PCR amplification of 50 ng of genomic DNA template. The primers were designed with Primer 3 (http://frodo.wi.mit.edu/primer3) and are detailed in Table S6. Bidirectional sequencing of the purified PCR products was performed by GATC Sequencing Facilities (Konstanz, Germany).

### RNA and protein analysis using the patient’s cells

Fibroblasts of patient II.4 and control individuals were obtained by skin biopsy as previously described.(Scheidecker et al., 2015) Three sex and age matched controls were used.

RNA was extracted from skin fibroblasts of individual II.4 and a healthy unrelated control using Rneasy RNA kit (Qiagen) then we performed reverse transcription using the iScriptTM cDNA Synthesis Kit (BioRad, Hercules, CA).

Protein analysis include Western Blot for which extracted proteins using the RIPA Buffer (89901 Thermo Scientific) complemented with protease inhibitor cocktail (Roche 06538282001) from primary fibroblasts of affected and control individuals were loaded onto Mini-PROTEAN TGX gels (BIO-RAD). Immunofluorescence assays were performed using primary fibroblasts from patient and control individuals grown in Nunc Lab-Tek chamber slides (Thermo Scientific, Waltham, MA, USA) fixed with 4% paraformaldehyde, incubated with PBS 0,5% Triton X-100 for 10min and blocked with PBS-20% FCS. Cells were then incubated for 1 h with primary antibodies, washed three times in PBS, incubated for 1 h with secondary antibody and DAPI, washed again in PBS and mounted in Elvanol No-Fade mounting medium before observation on a fluorescence microscope (Zeiss Axio Observer D1) at a X400 magnification. The primary antibodies were directed against the N- and C-terminal part of PSMC3. The secondary antibody was goat anti-mouse Alexa Fluor coupled 568 IgG (Invitrogen).

Accumulation of ubiquinated proteins has been observed using a dedicated western blot. Skin fibroblasts from three controls and the affected individual were recovered in ice cold RIPA buffer with protease inhibitors (‘Complete EDTA-free’; Roche Diagnostics, 1 tablet in 10ml buffer) and 25mM N-ethylmaleimide (NEM, diluted freshly in ethanol to prevent artifactual deubiquitination), left on ice for 45min, centrifuged for 10min at 12000rpm, supernatant recovered and 5X Laemmli buffer added. Bands were quantified relative to the total amount of protein loaded (stainfree) using Image Lab. All bands corresponding to the various ubiquitinated proteins were added and expressed relative to the amount of ubiquitinated proteins in control 1 fibroblasts. The mean of six independent experiments was calculated and represented as a histogram and a student test performed to determine the significance. Primary and secondary antibodies used in these experiments study as well as their dilution are described in Table S7.

### Co-immunoprecipitation and mass-spectrometry analysis

PSMC3 protein and its partners were immunoprecipitated from patient’s fibroblasts using magnetic microparticles (MACS purification system, Miltenyi Biotech) according to the manufacturer’s instructions and as previously described (Stoetzel et al., 2016, 15). Briefly, PSMC3 complexes were captured with an anti-PSMC3 antibody (Abcam ab171969) and the target and its associated proteins were purified on the protein G microbeads (Miltenyi Biotech). Proteins were eluted out of the magnetic stand with 1x laemmli buffer (Biorad). To optimize reproducibility, co-immunoprecipitation experiments were carried out in affinity triplicates on exactly 1.1 mg of proteins for each sample. For negative controls, halves of each sample were pooled and immunoprecipitated with the protein G beads, omitting the antibody.

Protein extracts were prepared as described in a previous study (Waltz et al., 2019). Each sample was precipitated with 0.1 M ammonium acetate in 100% methanol, and proteins were resuspended in 50 mM ammonium bicarbonate. After a reduction-alkylation step (dithiothreitol 5 mM – iodoacetamide 10 mM), proteins were digested overnight with sequencing-grade porcine trypsin (1:25, w/w, Promega, Fitchburg, MA, USA). The resulting vacuum-dried peptides were resuspended in water containing 0.1% (v/v) formic acid (solvent A). One fifth of the peptide mixtures were analyzed by nanoLC-MS/MS an Easy-nanoLC-1000 system coupled to a Q-Exactive Plus mass spectrometer (Thermo-Fisher Scientific, USA) operating in positive mode. Five microliters of each sample were loaded on a C-18 precolumn (75 μm ID × 20 mm nanoViper, 3µm Acclaim PepMap; Thermo) coupled with the analytical C18 analytical column (75 μm ID × 25 cm nanoViper, 3 µm Acclaim PepMap; Thermo). Peptides were eluted with a 160 min gradient of 0.1% formic acid in acetonitrile at 300 nL/min. The Q-Exactive Plus was operated in data-dependent acquisition mode (DDA) with Xcalibur software (Thermo-Fisher Scientific). Survey MS scans were acquired at a resolution of 70K at 200 m/z (mass range 350-1250), with a maximum injection time of 20 ms and an automatic gain control (AGC) set to 3e6. Up to 10 of the most intense multiply charged ions (≥2) were selected for fragmentation with a maximum injection time of 100 ms, an AGC set at 1e5 and a resolution of 17.5K. A dynamic exclusion time of 20 s was applied during the peak selection process.

MS data were searched against the Swissprot database (release 2019_05) with Human taxonomy. We used the Mascot algorithm (version 2.5, Matrix Science) to perform the database search with a decoy strategy and search parameters as follows: carbamidomethylation of cysteine, N-terminal protein acetylation and oxidation of methionine were set as variable modifications; tryptic specificity with up to three missed cleavages was used. The mass tolerances in MS and MS/MS were set to 10 ppm and 0.05 Da respectively, and the instrument configuration was specified as “ESI-Trap”. The resulting .dat Mascot files were then imported into Proline v1.4 package (http://proline.profiproteomics.fr/) for post-processing. Proteins were validated with Mascot pretty rank equal to 1, 1% FDR on both peptide spectrum matches (PSM) and protein sets (based on score). The total number of MS/MS fragmentation spectra was used to quantify each protein in the different samples.

For the statistical analysis of the co-immunoprecipitation data, we compared the data collected from multiple experiments against the negative control IPs using a homebrewed R package as described previsously (Chicois et al., 2018) except that that the size factor used to scale samples were calculated according to the DESeq normalisation method (i.e. median of ratios method (Anders and Huber, 2010)). The package calculates the fold change and an adjusted P-value corrected by Benjamini– Hochberg for each identified protein (and visualizes the data in volcano plots).

### Zebrafish analysis

#### Zebrafish (*Danio rerio*) maintenance and husbandry

In this study the zebrafish wild-type line AB2O2 (University of Oregon, Eugene) and the transgenic line gSAIzGFF539A (SOKENDAI, the Graduate University for Advanced Studies, Mishima, Japan) were used and maintained at 28°C under a 14 hour light and 10 hour dark cycle as described previously.(Westerfield, 2000) When fish reached sexual maturity, zebrafish couples were transferred to breeding tanks the day before and crossed after the beginning of the next light cycle. Fertilized zebrafish eggs were raised at 28.5 °C in 1× Instant Ocean salt solution (Aquarium Systems, Inc.). Zebrafish husbandry and experimental procedures were performed in accordance with German animal protection regulations (Regierungspräsidium, Karlsruhe, Germany, AZ35-9185.81/G-137/10).

#### Microinjections

Injections were performed as described before.(Müller et al., 1999) Morpholinos *psmc3-mo* (TGTGAATCACAGTATGAAGCGTGCC, Genetools LLC, Oregon) and *ctrl-mo* (5-bp mismatched) (TGT<u>C</u>AAT<u>G</u>A<u>G</u>AGTAT<u>C</u>AA<u>C</u>CGTGCC, Genetools LLC, Oregon) were injected at 0.2 mM (Figure S14). CRISPR guide RNAs were designed with ChopChop software and synthesized with the MEGAshortscript T7 Transcription Kit (Ambion) according to the manufacturer’s instructions. For CRISPR/Cas9 injections 300 ng/μl of Cas9 protein (GeneArt Platinum Cas9 Nuclease, Invitrogen) and 300 ng/µl guide RNA were combined. Guide RNA1 (G1 binding sequence: AGATCCTAATGACCAAGAGG<u>AGG</u>) targets exon 4 and Guide RNA2 (G2 binding sequence: CAGGATATCCACCCTGTTAG<u>TGG</u>) targets exon 9. The web interface PCR-F-Seq q (http://iai-gec-server.iai.kit.edu) was used to quantify the cutting efficiency of both guide RNAs.(Etard et al., 2017) For the life imaging experiment, the transgenic line gSAIzGFF539A, marking the semicircular canals, was generated using the Gal4-UAS system as described previously.(Asakawa et al., 2008) For rescue experiments, mRNA of zebrafish *psmc3* was synthesized using the mMESSAGE mMACHINE system (Ambion). Zebrafish *psmc3* mRNA was injected at a final concentration of 10 ng/µl.

#### PCR amplification and cloning

Total RNA from 24 to 72 hpf zebrafish embryos was extracted using Tri-reagent (Invitrogen, Carlsbad, CA) and reverse transcribed using M-MLV Reverse Transcriptase (Promega, Germany). To analyse the expression pattern of *psmc3* in zebrafish embryos we amplified and cloned a 956 bp *psmc3* fragment into the pGEMT-easy vector (Promega). For the synthesis of DIG-labelled RNA probes, we used Apa1 to linearize the plasmid and SP6 to transcribe the anti-sense RNA DIG probes. To exclude possible off-target effects caused by morpholino or CRISPR/Cas9 injection a rescue experiment was performed. Full-length *psmc3* was cloned into pCS2+ using EcoRI, linearized with NotI and *psmc3* full-length RNA synthesised using the mMESSAGE mMACHINE SP6 Kit (Ambion). To verify the efficiency of morpholino (*psmc3*-*mo*), we extracted total RNA from morpholino injected embryos using Tri-reagent (Invitrogen, Carlsbad, CA), transcribed mRNA into cDNA using M-MLV Reverse Transcriptase (Promega, Germany), and checked the effect on the *psmc3* splice sites by RT-PCR using the amplification conditions as described before (Figure S14B).(Müller et al., 1999) Using Sanger Sequencing, the CRISPR/Cas9 efficiency was assessed (Figure S14C-D).

#### Whole-mount *in situ* hybridization

In situ hybridization was performed as previously described.(Costa et al., 2002; Crow and Stockdale, 1986; Oxtoby and Jowett, 1993) To suppress melanogenesis, 20 hpf zebrafish embryos were transferred and raised in water supplemented with 200 mM 1-phenyl 2-thiourea (PTU). Probes targeting the messenger RNAs of the genes *krox20, msxc, her8a, sox19b* were obtained from Armant *et al.* (2013).(Armant et al., 2013) Plasmids to generate probes targeting *otopetrin, versican and versican b* were provided by the laboratory of Tanya Whitfield.

#### Immunohistochemistry

Immunostaining of whole zebrafish embryos after 5 dpf was performed as previously described(Leventea et al., 2016) and imaged with Leica TCS SP5 confocal microscope. Antibodies are indicated in Table S7.

#### Microscopy

To examine the eye and ear phenotype in morphants and crispants, we anaesthetized living embryos with 0.0168% (w/v) MESAB (tricane methanesulfonate, MS-222; Sigma-Aldrich, Taufkirchen, Germany) and embedded them into 0.5% (w/v) low melting point agarose chilled to 37°C in a lateral position with one eye/ear facing towards the objective. Confocal reflection microscopy was used to examine zebrafish eyes for abnormal light reflection evoked through cataract as previously described(Takamiya et al., 2016) and imaged with the Leica TCS SP2 confocal system with a 63x water immersion objective. Brightfield and fluorescence real time imaging of zebrafish ears were acquired using the Leica TCS SP5 confocal microscope with a 63x and 40x water immersion objective.

### Native-PAGE and proteasome in-Gel peptidase activity assay

Patient and control fibroblasts were lysed in equal amounts of TSDG buffer (10 mM Tris pH 7.0, 10 mM NaCl, 25 mM KCl, 1.1 mM MgCl_2_, 0.1 mM EDTA, 2 mM DTT, 2 mM ATP, 1 mM NaN_3_, 20 % Glycerol) prior to protein quantification using a standard BCA assay (Pierce) following the manufacturer’s instructions. Twenty micrograms of whole-cell extracts were loaded on 3-12% gradient native-PAGE gels (Invitrogen) and subsequently subjected to a 16-h electrophoresis at 45 V using a running buffer consisting of 50 mM BisTris/Tricine, pH 6,8. Chymotrypsin-like activity of the separated proteasome complexes was then measured by incubating 0.1mM of the suc-LLVY-AMC substrate (Bachem) at 37°C for 20 min in a final volume of 10 ml of overlay buffer (20 mM Tris, 5 mM MgCl_2_, pH 7,0. Proteasome bands were then visualised by exposure of the gel to UV light at 360 nm and detected at 460 nm using an Imager

### Measurement of proteasome activity in crude extracts

Ten micrograms of whole-cell extracts deriving from control and patient fibroblasts were tested for chymotrypsin-like activity by exposing the cell lysates to 0.1 mM of the Suc-LLVY-AMC peptide. Assays were carried out in a 100 µl reaction volume of ATP/DTT lysis buffer at 37°C. The rate of cleavage of fluorogenic peptide substrate was determined by monitoring the fluorescence of released aminomethylcoumarin using a plate reader at an excitation wavelength of 360 nm and emission wavelength of 460 nm over a period of 120 min.

### Western blot analysis

For western blotting, equal amounts of RIPA (50 mM Tris pH 7.5, 150 mM NaCl, 2 mM EDTA, 1 mM N-ethylmaleimide, 10 µM MG-132, 1% NP40, 0.1% SDS) buffered protein extracts from control and patient fibroblasts were separated in SDS-Laemmli gels (12,5 or 10%). Briefly, following separation, proteins were transferred to PVDF membranes (200V for 1 h) blocked with 1X Roti®-Block (Carl Roth) for 20 min at room temperature under shaking and subsequently probed overnight at 4°C with relevant primary antibodies. The anti-Alpha6 (clone MCP20), anti-Beta1 (clone MCP421), anti-Beta2 (clone MCP165), anti-Rpt1 (PSMC2, BML-PW8315), anti-Rpt2 (PSMC1, BML-PW0530), anti-Rpt3 (PSMC4, clone TBP7-27), anti-Rpt4 (PSMC6, clone p42-23), anti-Rpt5 (PSMC3, BML-PW8310) and anti-Rpt6 (PSMC5, clone p45-110) primary antibodies were purchased from Enzo Life Sciences. The anti-Beta1i (LMP2, K221) and anti-PA28-α (K232/1) are laboratory stocks and were used in previous studies(Poli et al., 2018). Antibodies specific for Beta5 (ab3330) and α-tubulin (clone DM1A, ab7291) were purchased from Abcam. Primary antibodies specific for TCF11/Nrf1 (clone D5B10) and ubiquitin (clone D9D5) were obtained from Cell Signaling Technology. The anti-Beta5i antibody (LMP7, clone A12) was a purchase from Santa Cruz Biotechnology Inc. After incubation with the primary antibodies, membranes were washed three times with PBS/0.4% Tween and incubated with anti-mouse or –rabbit HRP conjugated secondary antibodies (1/5.000). Visualisation of immunoreactive proteins was performed with enhanced chemiluminescence detection kit (ECL) (Biorad).

## AVAILABILITY OF DATA

Data generated or analyzed during this study are included in the published article and the corresponding supplementary data. The whole-genome sequencing data have been deposited at the European Genome-Phenome Archive (EGA), which is hosted by the European Bioinformatics Institute (EBI) and the Centre for Genomic Regulation (CRG), under accession number EGAS00001003942. The mass spectrometry proteomic data have been deposited at the PRIDE PRoteomics IDEntifications database (PRIDE) hosted by the EBI, under ProteomeXchange identifier PXD015836 and 10.6019/PXD015836. Access to this data possible to the reviewers using the following identifiers: username, password: iQ2HoDAx. *PSMC3* variants have been submitted to ClinVar with the following accession number: SCV000864220.1 (https://www.ncbi.nlm.nih.gov/clinvar). The rest of the data that support the findings of this study are available from the corresponding authors on reasonable request.

## CONFLICTS OF INTEREST

The authors declare no conflict of interest.

## ACKNOWLEDGMENTS

We would like to thank the patients and their family for their participation. Whole exome sequencing was performed by the IGBMC Microarray and Sequencing platform, a member of the “France Génomique” consortium (ANR-10-INBS-0009) and funded thanks to the support of the Centre Régional de Génétique Médicale de Strasbourg (CREGEMES). Whole genome sequencing was supported by the Laboratory of Excellence GENMED (Medical Genomics) grant no. ANR-10-LABX-0013 managed by the National Research Agency (ANR) part of the Investment for the Future (*Investissement d’Avenir*) program. This work was also supported by grants from the Molecular Medicine Research Consortium of the University of Greifswald (FOVB-2018-11 to FE), the National BioResource Project (NBRP) and the NBRP Fundamental Technologies Upgrading Program from AMED (Japan Agency for Medical Research and Development). A.K-H. is supported by a doctoral fellowship from the “Initiatives d’Excellence” (IdEx) through the University of Strasbourg and by the Franco-German University (UFA/DFH). S.F. and SB are supported by CNRS/Université de Strasbourg and S.B. also by INSERM. We wish to thank our colleagues around the world for testing their cohorts of patients (Elena Semina, Patrick Calvas, Sandrine Marlin, Alain Verloes). We thank Frederic Plewniak for the use of HomoSNP. We thank Tanja Whitfield for sharing plasmids for the examination of the zebrafish ear. The computing resources for this work were provided by the BICS and BISTRO bioinformatics platforms in Strasbourg. The mass spectrometry instrumentation was funded by the University of Strasbourg, IdEx “Equipement mi-lourd” 2015.

## AUTHOR CONTRIBUTIONS

E.S., S.S., F.S., V.P., C.SS., V.L., D.L. and H.D. gathered data from patients and performed clinical investigations; S.M., A.B., JF.D., I.P., V.G. and K.C. gathered sequencing data and performed analyses; A.KH., F.E., C.S., S.B, B.A.Z. and S.F. performed cell biology experiments and data analyses; C.S., L.K., J.C. and P.H. designed and performed the mass spectrometry experiments and data analyses; A.KH., M.T., K.K., C.E. and U.S. designed and performed the zebrafish experiments and data analyses; A. KH., J.M., U.W and H.D. analysed the data and wrote the paper; F.E., V.G., E.S., S.S., S.F, S.B., C.S., U.S., P.H., C.E. and EK contributed to manuscript writing; J.M., U.W. and H.D provided direction for the project, conceived and designed the experiments.

## REFERENCES

1000 Genomes Project Consortium, A. Auton, L.D. Brooks, R.M. Durbin, E.P. Garrison, H.M. Kang, J.O. Korbel, J.L. Marchini, S. McCarthy, G.A. McVean, and G.R. Abecasis. 2015. A global reference for human genetic variation. Nature. 526:68–74. doi:10.1038/nature15393.

Adzhubei, I.A., S. Schmidt, L. Peshkin, V.E. Ramensky, A. Gerasimova, P. Bork, A.S. Kondrashov, and S.R. Sunyaev. 2010. A method and server for predicting damaging missense mutations. Nat. Methods. 7:248–249. doi:10.1038/nmeth0410-248.

Agarwal, A.K., C. Xing, G.N. DeMartino, D. Mizrachi, M.D. Hernandez, A.B. Sousa, L. Martinez de Villarreal, H.G. dos Santos, and A. Garg. 2010. PSMB8 encoding the beta5i proteasome subunit is mutated in joint contractures, muscle atrophy, microcytic anemia, and panniculitis-induced lipodystrophy syndrome. Am J Hum Genet. 87:866–72. doi:10.1016/j.ajhg.2010.10.031.

Anders, S., and W. Huber. 2010. Differential expression analysis for sequence count data. Genome Biol. 11:R106. doi:10.1186/gb-2010-11-10-r106.

Arima, K., A. Kinoshita, H. Mishima, N. Kanazawa, T. Kaneko, T. Mizushima, K. Ichinose, H. Nakamura, A. Tsujino, A. Kawakami, M. Matsunaka, S. Kasagi, S. Kawano, S. Kumagai, K. Ohmura, T. Mimori, M. Hirano, S. Ueno, K. Tanaka, M. Tanaka, I. Toyoshima, H. Sugino, A. Yamakawa, K. Tanaka, N. Niikawa, F. Furukawa, S. Murata, K. Eguchi, H. Ida, and K. Yoshiura. 2011. Proteasome assembly defect due to a proteasome subunit beta type 8 (PSMB8) mutation causes the autoinflammatory disorder, Nakajo-Nishimura syndrome. Proc Natl Acad Sci U A. 108:14914–9. doi:10.1073/pnas.1106015108.

Armant, O., M. März, R. Schmidt, M. Ferg, N. Diotel, R. Ertzer, J.C. Bryne, L. Yang, I. Baader, M. Reischl, J. Legradi, R. Mikut, D. Stemple, W. van IJcken, A. van der Sloot, B. Lenhard, U. Strähle, and S. Rastegar. 2013. Genome-wide, whole mount in situ analysis of transcriptional regulators in zebrafish embryos. Dev. Biol. 380:351–362. doi:10.1016/j.ydbio.2013.05.006.

Asakawa, K., M.L. Suster, K. Mizusawa, S. Nagayoshi, T. Kotani, A. Urasaki, Y. Kishimoto, M. Hibi, and K. Kawakami. 2008. Genetic dissection of neural circuits by Tol2 transposon-mediated Gal4 gene and enhancer trapping in zebrafish. Proc. Natl. Acad. Sci. 105:1255–1260. doi:10.1073/pnas.0704963105.

Azaiez, H., K.T. Booth, S.S. Ephraim, B. Crone, E.A. Black-Ziegelbein, R.J. Marini, A.E. Shearer, C.M. Sloan-Heggen, D. Kolbe, T. Casavant, M.J. Schnieders, C. Nishimura, T. Braun, and R.J.H. Smith. 2018. Genomic Landscape and Mutational Signatures of Deafness-Associated Genes. Am. J. Hum. Genet. 103:484–497. doi:10.1016/j.ajhg.2018.08.006.

Backenroth, D., J. Homsy, L.R. Murillo, J. Glessner, E. Lin, M. Brueckner, R. Lifton, E. Goldmuntz, W.K. Chung, and Y. Shen. 2014. CANOES: detecting rare copy number variants from whole exome sequencing data. Nucleic Acids Res. 42:e97. doi:10.1093/nar/gku345.

Bartenhagen, C., and M. Dugas. 2016. Robust and exact structural variation detection with paired-end and soft-clipped alignments: SoftSV compared with eight algorithms. Brief. Bioinform. 17:51–62. doi:10.1093/bib/bbv028.

Belkadi, A., A. Bolze, Y. Itan, A. Cobat, Q.B. Vincent, A. Antipenko, L. Shang, B. Boisson, J.-L. Casanova, and L. Abel. 2015. Whole-genome sequencing is more powerful than whole-exome sequencing for detecting exome variants. Proc. Natl. Acad. Sci. U. S. A. 112:5473–5478. doi:10.1073/pnas.1418631112.

Brehm, A., Y. Liu, A. Sheikh, B. Marrero, E. Omoyinmi, Q. Zhou, G. Montealegre, A. Biancotto, A. Reinhardt, A. Almeida de Jesus, M. Pelletier, W.L. Tsai, E.F. Remmers, L. Kardava, S. Hill, H. Kim, H.J. Lachmann, A. Megarbane, J.J. Chae, J. Brady, R.D. Castillo, D. Brown, A.V. Casano, L. Gao, D. Chapelle, Y. Huang, D. Stone, Y. Chen, F. Sotzny, C.C. Lee, D.L. Kastner, A. Torrelo, A. Zlotogorski, S. Moir, M. Gadina, P. McCoy, R. Wesley, K.I. Rother, P.W. Hildebrand, P. Brogan, E. Kruger, I. Aksentijevich, and R. Goldbach-Mansky. 2015. Additive loss-of-function proteasome subunit mutations in CANDLE/PRAAS patients promote type I IFN production. J Clin Invest. 125:4196–211. doi:10.1172/JCI81260.

Busch-Nentwich, E. 2004. The deafness gene dfna5 is crucial for ugdh expression and HA production in the developing ear in zebrafish. Development. 131:943–951. doi:10.1242/dev.00961.

Chen, S., J. Wu, Y. Lu, Y.-B. Ma, B.-H. Lee, Z. Yu, Q. Ouyang, D.J. Finley, M.W. Kirschner, and Y. Mao. 2016. Structural basis for dynamic regulation of the human 26S proteasome. Proc. Natl. Acad. Sci. 113:12991–12996. doi:10.1073/pnas.1614614113.

Chicois, C., H. Scheer, S. Garcia, H. Zuber, J. Mutterer, J. Chicher, P. Hammann, D. Gagliardi, and D. Garcia. 2018. The UPF1 interactome reveals interaction networks between RNA degradation and translation repression factors in Arabidopsis. Plant J. Cell Mol. Biol. 96:119–132. doi:10.1111/tpj.14022.

Costa, M.L., R.C. Escaleira, V.B. Rodrigues, M. Manasfi, and C.S. Mermelstein. 2002. Some distinctive features of zebrafish myogenesis based on unexpected distributions of the muscle cytoskeletal proteins actin, myosin, desmin, alpha-actinin, troponin and titin. Mech. Dev. 116:95–104.

Crow, M.T., and F.E. Stockdale. 1986. Myosin expression and specialization among the earliest muscle fibers of the developing avian limb. Dev. Biol. 113:238–254.

Cruz, S., J.-C. Shiao, B.-K. Liao, C.-J. Huang, and P.-P. Hwang. 2009. Plasma membrane calcium ATPase required for semicircular canal formation and otolith growth in the zebrafish inner ear. J. Exp. Biol. 212:639–647. doi:10.1242/jeb.022798.

Cvekl, A., and X. Zhang. 2017. Signaling and Gene Regulatory Networks in Mammalian Lens Development. Trends Genet. 33:677–702. doi:10.1016/j.tig.2017.08.001.

De Franco, E., S.E. Flanagan, T. Yagi, D. Abreu, J. Mahadevan, M.B. Johnson, G. Jones, F. Acosta, M. Mulaudzi, N. Lek, V. Oh, O. Petz, R. Caswell, S. Ellard, F. Urano, and A.T. Hattersley. 2017. Dominant ER Stress–Inducing *WFS1* Mutations Underlie a Genetic Syndrome of Neonatal/Infancy-Onset Diabetes, Congenital Sensorineural Deafness, and Congenital Cataracts. Diabetes. 66:2044–2053. doi:10.2337/db16-1296.

DePristo, M.A., E. Banks, R. Poplin, K.V. Garimella, J.R. Maguire, C. Hartl, A.A. Philippakis, G. del Angel, M.A. Rivas, M. Hanna, A. McKenna, T.J. Fennell, A.M. Kernytsky, A.Y. Sivachenko, K. Cibulskis, S.B. Gabriel, D. Altshuler, and M.J. Daly. 2011. A framework for variation discovery and genotyping using next-generation DNA sequencing data. Nat. Genet. 43:491–498. doi:10.1038/ng.806.

Deveraux, Q., V. Ustrell, C. Pickart, and M. Rechsteiner. 1994. A 26 S protease subunit that binds ubiquitin conjugates. J. Biol. Chem. 269:7059–7061.

Elsen, G.E., L.Y. Choi, V.E. Prince, and R.K. Ho. 2009. The autism susceptibility gene met regulates zebrafish cerebellar development and facial motor neuron migration. Dev. Biol. 335:78–92. doi:10.1016/j.ydbio.2009.08.024.

Etard, C., S. Joshi, J. Stegmaier, R. Mikut, and U. Strähle. 2017. Tracking of Indels by DEcomposition is a Simple and Effective Method to Assess Efficiency of Guide RNAs in Zebrafish. Zebrafish. 14:586–588. doi:10.1089/zeb.2017.1454.

Froyen, G., M. Corbett, J. Vandewalle, I. Jarvela, O. Lawrence, C. Meldrum, M. Bauters, K. Govaerts, L. Vandeleur, H. Van Esch, J. Chelly, D. Sanlaville, H. van Bokhoven, H.-H. Ropers, F. Laumonnier, E. Ranieri, C.E. Schwartz, F. Abidi, P.S. Tarpey, P.A. Futreal, A. Whibley, F.L. Raymond, M.R. Stratton, J.-P. Fryns, R. Scott, M. Peippo, M. Sipponen, M. Partington, D. Mowat, M. Field, A. Hackett, P. Marynen, G. Turner, and J. Gécz. 2008. Submicroscopic Duplications of the Hydroxysteroid Dehydrogenase HSD17B10 and the E3 Ubiquitin Ligase HUWE1 Are Associated with Mental Retardation. Am. J. Hum. Genet. 82:432–443. doi:10.1016/j.ajhg.2007.11.002.

Gao, M., Y. Huang, L. Wang, M. Huang, F. Liu, S. Liao, S. Yu, Z. Lu, S. Han, X. Hu, Z. Qu, X. Liu, T. Assefa Yimer, L. Yang, Z. Tang, D.W.-C. Li, and M. Liu. 2017. HSF4 regulates lens fiber cell differentiation by activating p53 and its downstream regulators. Cell Death Dis. 8:e3082. doi:10.1038/cddis.2017.478.

Geng, F.-S., L. Abbas, S. Baxendale, C.J. Holdsworth, A.G. Swanson, K. Slanchev, M. Hammerschmidt, J. Topczewski, and T.T. Whitfield. 2013. Semicircular canal morphogenesis in the zebrafish inner ear requires the function of gpr126 (lauscher), an adhesion class G protein-coupled receptor gene. Development. 140:4362–4374. doi:10.1242/dev.098061.

Geoffroy, V., Y. Herenger, A. Kress, C. Stoetzel, A. Piton, H. Dollfus, and J. Muller. 2018a. AnnotSV: An integrated tool for Structural Variations annotation. Bioinforma. Oxf. Engl. doi:10.1093/bioinformatics/bty304.

Geoffroy, V., C. Pizot, C. Redin, A. Piton, N. Vasli, C. Stoetzel, A. Blavier, J. Laporte, and J. Muller. 2015. VaRank: a simple and powerful tool for ranking genetic variants. PeerJ. 3:e796. doi:10.7717/peerj.796.

Geoffroy, V., C. Stoetzel, S. Scheidecker, E. Schaefer, I. Perrault, S. Bär, A. Kröll, M. Delbarre, M. Antin, A.-S. Leuvrey, C. Henry, H. Blanché, E. Decker, K. Kloth, G. Klaus, C. Mache, D. Martin-Coignard, S. McGinn, A. Boland, J.-F. Deleuze, S. Friant, S. Saunier, J.-M. Rozet, C. Bergmann, H. Dollfus, and J. Muller. 2018b. Whole-genome sequencing in patients with ciliopathies uncovers a novel recurrent tandem duplication in IFT140. Hum. Mutat. 39:983–992. doi:10.1002/humu.23539.

Haddon, C.M., and J.H. Lewis. 1991. Hyaluronan as a propellant for epithelial movement: the development of semicircular canals in the inner ear of Xenopus. Dev. Camb. Engl. 112:541–550.

Han, Y., Y. Mu, X. Li, P. Xu, J. Tong, Z. Liu, T. Ma, G. Zeng, S. Yang, J. Du, and A. Meng. 2011. Grhl2 deficiency impairs otic development and hearing ability in a zebrafish model of the progressive dominant hearing loss DFNA28. Hum. Mol. Genet. 20:3213–3226. doi:10.1093/hmg/ddr234.

Huang, N., I. Lee, E.M. Marcotte, and M.E. Hurles. 2010. Characterising and Predicting Haploinsufficiency in the Human Genome. PLOS Genet. 6:e1001154. doi:10.1371/journal.pgen.1001154.

Husnjak, K., S. Elsasser, N. Zhang, X. Chen, L. Randles, Y. Shi, K. Hofmann, K.J. Walters, D. Finley, and I. Dikic. 2008. Proteasome subunit Rpn13 is a novel ubiquitin receptor. Nature. 453:481–488. doi:10.1038/nature06926.

Imai, F., A. Yoshizawa, N. Fujimori-Tonou, K. Kawakami, and I. Masai. 2010. The ubiquitin proteasome system is required for cell proliferation of the lens epithelium and for differentiation of lens fiber cells in zebrafish. Development. 137:3257–3268. doi:10.1242/dev.053124.

Karczewski, K.J., L.C. Francioli, G. Tiao, B.B. Cummings, J. Alföldi, Q. Wang, R.L. Collins, K.M. Laricchia,A. Ganna, D.P. Birnbaum, L.D. Gauthier, H. Brand, M. Solomonson, N.A. Watts, D. Rhodes, M. Singer-Berk, E.G. Seaby, J.A. Kosmicki, R.K. Walters, K. Tashman, Y. Farjoun, E. Banks, T. Poterba, A. Wang, C. Seed, N. Whiffin, J.X. Chong, K.E. Samocha, E. Pierce-Hoffman, Z. Zappala, A.H. O’Donnell-Luria, E. Vallabh Minikel, B. Weisburd, M. Lek, J.S. Ware, C. Vittal, I.M. Armean,L. Bergelson, K. Cibulskis, K.M. Connolly, M. Covarrubias, S. Donnelly, S. Ferriera, S. Gabriel, J. Gentry, N. Gupta, T. Jeandet, D. Kaplan, C. Llanwarne, R. Munshi, S. Novod, N. Petrillo, D. Roazen, V. Ruano-Rubio, A. Saltzman, M. Schleicher, J. Soto, K. Tibbetts, C. Tolonen, G. Wade,M.E. Talkowski, B.M. Neale, M.J. Daly, and D.G. MacArthur. 2019. Variation across 141,456 human exomes and genomes reveals the spectrum of loss-of-function intolerance across human protein-coding genes. bioRxiv. 531210. doi:10.1101/531210.

Khalil, R., C. Kenny, R.S. Hill, G.H. Mochida, R. Nasir, J.N. Partlow, B.J. Barry, M. Al-Saffar, C. Egan, C.R. Stevens, S.B. Gabriel, A.J. Barkovich, J.W. Ellison, L. Al-Gazali, C.A. Walsh, and M.H. Chahrour. 2018. *PSMD12* haploinsufficiency in a neurodevelopmental disorder with autistic features. Am.J. Med. Genet. B Neuropsychiatr. Genet. 177:736–745. doi:10.1002/ajmg.b.32688.

Kindt, K.S., G. Finch, and T. Nicolson. 2012. Kinocilia Mediate Mechanosensitivity in Developing Zebrafish Hair Cells. Dev. Cell. 23:329–341. doi:10.1016/j.devcel.2012.05.022.

Kishino, T., M. Lalande, and J. Wagstaff. 1997. UBE3A/E6-AP mutations cause Angelman syndrome. Nat. Genet. 15:70–73. doi:10.1038/ng0197-70.

Küry, S., T. Besnard, F. Ebstein, T.N. Khan, T. Gambin, J. Douglas, C.A. Bacino, W.J. Craigen, S.J. Sanders,A. Lehmann, X. Latypova, K. Khan, M. Pacault, S. Sacharow, K. Glaser, E. Bieth, L. Perrin-Sabourin, M.-L. Jacquemont, M.T. Cho, E. Roeder, A.-S. Denommé-Pichon, K.G. Monaghan, B. Yuan, F. Xia, S. Simon, D. Bonneau, P. Parent, B. Gilbert-Dussardier, S. Odent, A. Toutain, L. Pasquier, D. Barbouth, C.A. Shaw, A. Patel, J.L. Smith, W. Bi, S. Schmitt, W. Deb, M. Nizon, S. Mercier, M. Vincent, C. Rooryck, V. Malan, I. Briceño, A. Gómez, K.M. Nugent, J.B. Gibson, B. Cogné, J.R. Lupski, H.A.F. Stessman, E.E. Eichler, K. Retterer, Y. Yang, R. Redon, N. Katsanis, J.A. Rosenfeld, P.-M. Kloetzel, C. Golzio, S. Bézieau, P. Stankiewicz, and B. Isidor. 2017. De Novo Disruption of the Proteasome Regulatory Subunit PSMD12 Causes a Syndromic Neurodevelopmental Disorder. Am. J. Hum. Genet. 100:352–363. doi:10.1016/j.ajhg.2017.01.003.

Lam, Y.A., T.G. Lawson, M. Velayutham, J.L. Zweier, and C.M. Pickart. 2002. A proteasomal ATPase subunit recognizes the polyubiquitin degradation signal. Nature. 416:763–767. doi:10.1038/416763a.

Lek, M., K.J. Karczewski, E.V. Minikel, K.E. Samocha, E. Banks, T. Fennell, A.H. O’Donnell-Luria, J.S. Ware, A.J. Hill, B.B. Cummings, T. Tukiainen, D.P. Birnbaum, J.A. Kosmicki, L.E. Duncan, K. Estrada, F. Zhao, J. Zou, E. Pierce-Hoffman, J. Berghout, D.N. Cooper, N. Deflaux, M. DePristo,R. Do, J. Flannick, M. Fromer, L. Gauthier, J. Goldstein, N. Gupta, D. Howrigan, A. Kiezun, M.I. Kurki, A.L. Moonshine, P. Natarajan, L. Orozco, G.M. Peloso, R. Poplin, M.A. Rivas, V. Ruano-Rubio, S.A. Rose, D.M. Ruderfer, K. Shakir, P.D. Stenson, C. Stevens, B.P. Thomas, G. Tiao, M.T. Tusie-Luna, B. Weisburd, H.-H. Won, D. Yu, D.M. Altshuler, D. Ardissino, M. Boehnke, J. Danesh, S. Donnelly, R. Elosua, J.C. Florez, S.B. Gabriel, G. Getz, S.J. Glatt, C.M. Hultman, S. Kathiresan, M. Laakso, S. McCarroll, M.I. McCarthy, D. McGovern, R. McPherson, B.M. Neale, A. Palotie,S.M. Purcell, D. Saleheen, J.M. Scharf, P. Sklar, P.F. Sullivan, J. Tuomilehto, M.T. Tsuang, H.C. Watkins, J.G. Wilson, M.J. Daly, D.G. MacArthur, and Exome Aggregation Consortium. 2016. Analysis of protein-coding genetic variation in 60,706 humans. Nature. 536:285–291. doi:10.1038/nature19057.

Leventea, E., K. Hazime, C. Zhao, and J. Malicki. 2016. Analysis of cilia structure and function in zebrafish. In Methods in Cell Biology. Elsevier. 179–227.

Li, H., and R. Durbin. 2009. Fast and accurate short read alignment with Burrows-Wheeler transform. Bioinforma. Oxf. Engl. 25:1754–1760. doi:10.1093/bioinformatics/btp324.

Li, H., B. Handsaker, A. Wysoker, T. Fennell, J. Ruan, N. Homer, G. Marth, G. Abecasis, R. Durbin, and 1000 Genome Project Data Processing Subgroup. 2009. The Sequence Alignment/Map format and SAMtools. Bioinforma. Oxf. Engl. 25:2078–2079. doi:10.1093/bioinformatics/btp352.

Liu, Y., Y. Ramot, A. Torrelo, A.S. Paller, N. Si, S. Babay, P.W. Kim, A. Sheikh, C.C. Lee, Y. Chen, A. Vera,X. Zhang, R. Goldbach-Mansky, and A. Zlotogorski. 2012. Mutations in proteasome subunit beta type 8 cause chronic atypical neutrophilic dermatosis with lipodystrophy and elevated temperature with evidence of genetic and phenotypic heterogeneity. Arthritis Rheum. 64:895–907. doi:10.1002/art.33368.

Ma, C.P., C.A. Slaughter, and G.N. DeMartino. 1992. Identification, purification, and characterization of a protein activator (PA28) of the 20 S proteasome (macropain). J. Biol. Chem. 267:10515–10523.

MacDonald, J.R., R. Ziman, R.K.C. Yuen, L. Feuk, and S.W. Scherer. 2014. The Database of Genomic Variants: a curated collection of structural variation in the human genome. Nucleic Acids Res. 42:D986–D992. doi:10.1093/nar/gkt958.

Millimaki, B.B., E.M. Sweet, M.S. Dhason, and B.B. Riley. 2007. Zebrafish atoh1 genes: classic proneural activity in the inner ear and regulation by Fgf and Notch. Development. 134:295–305. doi:10.1242/dev.02734.

Mishra, S., S.-Y. Wu, A.W. Fuller, Z. Wang, K.L. Rose, K.L. Schey, and H.S. Mchaourab. 2018. Loss of αBcrystallin function in zebrafish reveals critical roles in the development of the lens and stress resistance of the heart. J. Biol. Chem. 293:740–753. doi:10.1074/jbc.M117.808634.

Moortgat, S., S. Berland, I. Aukrust, I. Maystadt, L. Baker, V. Benoit, A. Caro-Llopis, N.S. Cooper, F.-G. Debray, L. Faivre, T. Gardeitchik, B.I. Haukanes, G. Houge, E. Kivuva, F. Martinez, S.G. Mehta, M.-C. Nassogne, N. Powell-Hamilton, R. Pfundt, M. Rosello, T. Prescott, P. Vasudevan, B. van Loon, C. Verellen-Dumoulin, A. Verloes, C. von der Lippe, E. Wakeling, A.O.M. Wilkie, L. Wilson, A. Yuen, D. Study, K.J. Low, and R.A. Newbury-Ecob. 2018. HUWE1 variants cause dominant X-linked intellectual disability: a clinical study of 21 patients. Eur. J. Hum. Genet. EJHG. 26:64–74. doi:10.1038/s41431-017-0038-6.

Müller, F., P. Blader, S. Rastegar, N. Fischer, W. Knöchel, and U. Strähle. 1999. Characterization of zebrafish smad1, smad2 and smad5: the amino-terminus of smad1 and smad5 is required for specific function in the embryo. Mech. Dev. 88:73–88.

Nascimento, R.M.P., P.A. Otto, A.P.M. de Brouwer, and A.M. Vianna-Morgante. 2006. UBE2A, Which Encodes a Ubiquitin-Conjugating Enzyme, Is Mutated in a Novel X-Linked Mental Retardation Syndrome. Am. J. Hum. Genet. 79:549–555. doi:10.1086/507047.

Ng, P.C., and S. Henikoff. 2003. SIFT: Predicting amino acid changes that affect protein function. Nucleic Acids Res. 31:3812–3814.

Niceta, M., E. Stellacci, K.W. Gripp, G. Zampino, M. Kousi, M. Anselmi, A. Traversa, A. Ciolfi, D. Stabley,A. Bruselles, V. Caputo, S. Cecchetti, S. Prudente, M.T. Fiorenza, C. Boitani, N. Philip, D. Niyazov,C. Leoni, T. Nakane, K. Keppler-Noreuil, S.R. Braddock, G. Gillessen-Kaesbach, A. Palleschi, P.M. Campeau, B.H.L. Lee, C. Pouponnot, L. Stella, G. Bocchinfuso, N. Katsanis, K. Sol-Church, and M. Tartaglia. 2015. Mutations Impairing GSK3-Mediated MAF Phosphorylation Cause Cataract, Deafness, Intellectual Disability, Seizures, and a Down Syndrome-like Facies. Am. J. Hum. Genet. 96:816–825. doi:10.1016/j.ajhg.2015.03.001.

O’Leary, N.A., M.W. Wright, J.R. Brister, S. Ciufo, D. Haddad, R. McVeigh, B. Rajput, B. Robbertse, B. Smith-White, D. Ako-Adjei, A. Astashyn, A. Badretdin, Y. Bao, O. Blinkova, V. Brover, V. Chetvernin, J. Choi, E. Cox, O. Ermolaeva, C.M. Farrell, T. Goldfarb, T. Gupta, D. Haft, E. Hatcher, W. Hlavina, V.S. Joardar, V.K. Kodali, W. Li, D. Maglott, P. Masterson, K.M. McGarvey, M.R. Murphy, K. O’Neill, S. Pujar, S.H. Rangwala, D. Rausch, L.D. Riddick, C. Schoch, A. Shkeda, S.S. Storz, H. Sun, F. Thibaud-Nissen, I. Tolstoy, R.E. Tully, A.R. Vatsan, C. Wallin, D. Webb, W. Wu, M.J. Landrum, A. Kimchi, T. Tatusova, M. DiCuccio, P. Kitts, T.D. Murphy, and K.D. Pruitt. 2016. Reference sequence (RefSeq) database at NCBI: current status, taxonomic expansion, and functional annotation. Nucleic Acids Res. 44:D733–745. doi:10.1093/nar/gkv1189.

Oxtoby, E., and T. Jowett. 1993. Cloning of the zebrafish krox-20 gene (krx-20) and its expression during hindbrain development. Nucleic Acids Res. 21:1087–1095.

Paone, C., S. Rudeck, C. Etard, U. Strähle, W. Rottbauer, and S. Just. 2018. Loss of zebrafish Smyd1a interferes with myofibrillar integrity without triggering the misfolded myosin response. Biochem. Biophys. Res. Commun. 496:339–345. doi:10.1016/j.bbrc.2018.01.060.

Poli, M.C., F. Ebstein, S.K. Nicholas, M.M. de Guzman, L.R. Forbes, I.K. Chinn, E.M. Mace, T.P. Vogel, A.F. Carisey, F. Benavides, Z.H. Coban-Akdemir, R.A. Gibbs, S.N. Jhangiani, D.M. Muzny, C.M.B. Carvalho, D.A. Schady, M. Jain, J.A. Rosenfeld, L. Emrick, R.A. Lewis, B. Lee, members Undiagnosed Diseases Network, B.A. Zieba, S. Kury, E. Kruger, J.R. Lupski, B.L. Bostwick, and J.S. Orange. 2018. Heterozygous Truncating Variants in POMP Escape Nonsense-Mediated Decay and Cause a Unique Immune Dysregulatory Syndrome. Am J Hum Genet. 102:1126–1142. doi:10.1016/j.ajhg.2018.04.010.

Radhakrishnan, S.K., C.S. Lee, P. Young, A. Beskow, J.Y. Chan, and R.J. Deshaies. 2010. Transcription factor Nrf1 mediates the proteasome recovery pathway after proteasome inhibition in mammalian cells. Mol Cell. 38:17–28. doi:10.1016/j.molcel.2010.02.029.

Reese, M.G., F.H. Eeckman, D. Kulp, and D. Haussler. 1997. Improved splice site detection in Genie. J. Comput. Biol. J. Comput. Mol. Cell Biol. 4:311–323. doi:10.1089/cmb.1997.4.311.

Reis, L.M., and E.V. Semina. 2018. Genetic landscape of isolated pediatric cataracts: extreme heterogeneity and variable inheritance patterns within genes. Hum. Genet. doi:10.1007/s00439-018-1932-x.

Santiago-Sim, T., L.C. Burrage, F. Ebstein, M.J. Tokita, M. Miller, W. Bi, A.A. Braxton, J.A. Rosenfeld, M. Shahrour, A. Lehmann, B. Cogné, S. Küry, T. Besnard, B. Isidor, S. Bézieau, I. Hazart, H. Nagakura, L.L. Immken, R.O. Littlejohn, E. Roeder, Z. Afawi, R. Balling, N. Barisic, S. Baulac, D. Craiu, P. De Jonghe, R. Guerrero-Lopez, R. Guerrini, I. Helbig, H. Hjalgrim, J. Jähn, K.M. Klein, E. Leguern, H. Lerche, C. Marini, H. Muhle, F. Rosenow, J. Serratosa, K. Sterbová, A. Suls, R.S. Moller, P. Striano, Y. Weber, F. Zara, B. Kara, K. Hardies, S. Weckhuysen, P. May, J.R. Lemke, O. Elpeleg, B. Abu-Libdeh, K.N. James, J.L. Silhavy, M.Y. Issa, M.S. Zaki, J.G. Gleeson, J.R. Seavitt, M.E. Dickinson, M.C. Ljungberg, S. Wells, S.J. Johnson, L. Teboul, C.M. Eng, Y. Yang, P.-M. Kloetzel, J.D. Heaney, and M.A. Walkiewicz. 2017. Biallelic Variants in OTUD6B Cause an Intellectual Disability Syndrome Associated with Seizures and Dysmorphic Features. Am. J. Hum. Genet. 100:676–688. doi:10.1016/j.ajhg.2017.03.001.

Scheidecker, S., C. Etard, L. Haren, C. Stoetzel, S. Hull, G. Arno, V. Plagnol, S. Drunat, S. Passemard, A. Toutain, C. Obringer, M. Koob, V. Geoffroy, V. Marion, U. Strähle, P. Ostergaard, A. Verloes, A. Merdes, A.T. Moore, and H. Dollfus. 2015. Mutations in TUBGCP4 Alter Microtubule Organization via the γ-Tubulin Ring Complex in Autosomal-Recessive Microcephaly with Chorioretinopathy. Am. J. Hum. Genet. 96:666–674. doi:10.1016/j.ajhg.2015.02.011.

Shang, F., and A. Taylor. 2012. Chapter 10 - Role of the Ubiquitin–Proteasome in Protein Quality Control and Signaling: Implication in the Pathogenesis of Eye Diseases. In Progress in Molecular Biology and Translational Science. T. Grune, editor. Academic Press. 347–396.

Shapiro, M.B., and P. Senapathy. 1987. RNA splice junctions of different classes of eukaryotes: sequence statistics and functional implications in gene expression. Nucleic Acids Res. 15:7155–7174.

Singh, C.R., S. Lovell, N. Mehzabeen, W.Q. Chowdhury, E.S. Geanes, K.P. Battaile, and J. Roelofs. 2014. 1.15\AA resolution structure of the proteasome-assembly chaperone Nas2 PDZ domain. Acta Crystallogr. Sect. F. 70:418–423. doi:10.1107/S2053230X14003884.

Smith, D.M., S.-C. Chang, S. Park, D. Finley, Y. Cheng, and A.L. Goldberg. 2007. Docking of the Proteasomal ATPases’ Carboxyl Termini in the 20S Proteasome’s α Ring Opens the Gate for Substrate Entry. Mol. Cell. 27:731–744. doi:https://doi.org/10.1016/j.molcel.2007.06.033.

Sobreira, N., F. Schiettecatte, D. Valle, and A. Hamosh. 2015. GeneMatcher: A Matching Tool for Connecting Investigators with an Interest in the Same Gene. Hum. Mutat. 36:928–930. doi:10.1002/humu.22844.

Sotzny, F., E. Schormann, I. Kuhlewindt, A. Koch, A. Brehm, R. Goldbach-Mansky, K.E. Gilling, and E. Kruger. 2016. TCF11/Nrf1-Mediated Induction of Proteasome Expression Prevents Cytotoxicity by Rotenone. Antioxid Redox Signal. 25:870–885. doi:10.1089/ars.2015.6539.

Steffen, J., M. Seeger, A. Koch, and E. Kruger. 2010. Proteasomal degradation is transcriptionally controlled by TCF11 via an ERAD-dependent feedback loop. Mol Cell. 40:147–58. doi:10.1016/j.molcel.2010.09.012.

Stoetzel, C., S. Bär, J.-O. De Craene, S. Scheidecker, C. Etard, J. Chicher, J.R. Reck, I. Perrault, V. Geoffroy, K. Chennen, U. Strähle, P. Hammann, S. Friant, and H. Dollfus. 2016. A mutation in VPS15 (PIK3R4) causes a ciliopathy and affects IFT20 release from the cis-Golgi. Nat. Commun. 7:13586.

Stooke-Vaughan, G.A., N.D. Obholzer, S. Baxendale, S.G. Megason, and T.T. Whitfield. 2015. Otolith tethering in the zebrafish otic vesicle requires Otogelin and -Tectorin. Development. 142:1137–1145. doi:10.1242/dev.116632.

Takamiya, M., F. Xu, H. Suhonen, V. Gourain, L. Yang, N.Y. Ho, L. Helfen, A. Schröck, C. Etard, C. Grabher, S. Rastegar, G. Schlunck, T. Reinhard, T. Baumbach, and U. Strähle. 2016. Melanosomes in pigmented epithelia maintain eye lens transparency during zebrafish embryonic development. Sci. Rep. 6. doi:10.1038/srep25046.

Tanaka, K. 2009. The proteasome: Overview of structure and functions. Proc. Jpn. Acad. Ser. B. 85:12–36. doi:10.2183/pjab.85.12.

Teboul, L., S.A. Murray, and P.M. Nolan. 2017. Phenotyping first-generation genome editing mutants: a new standard? Mamm. Genome. 28:377–382. doi:10.1007/s00335-017-9711-x.

Thisse, B., and C. Thisse. 2004. Fast Release Clones: A High Throughput Expression Analysis. ZFIN Direct Data Submiss. Unpubl.

Thorvaldsdóttir, H., J.T. Robinson, and J.P. Mesirov. 2013. Integrative Genomics Viewer (IGV): high-performance genomics data visualization and exploration. Brief. Bioinform. 14:178–192. doi:10.1093/bib/bbs017.

Tyner, C., G.P. Barber, J. Casper, H. Clawson, M. Diekhans, C. Eisenhart, C.M. Fischer, D. Gibson, J.N. Gonzalez, L. Guruvadoo, M. Haeussler, S. Heitner, A.S. Hinrichs, D. Karolchik, B.T. Lee, C.M. Lee, P. Nejad, B.J. Raney, K.R. Rosenbloom, M.L. Speir, C. Villarreal, J. Vivian, A.S. Zweig, D. Haussler, R.M. Kuhn, and W.J. Kent. 2017. The UCSC Genome Browser database: 2017 update. Nucleic Acids Res. 45:D626–D634. doi:10.1093/nar/gkw1134.

Vaz-Drago, R., N. Custódio, and M. Carmo-Fonseca. 2017. Deep intronic mutations and human disease. Hum. Genet. 136:1093–1111. doi:10.1007/s00439-017-1809-4.

Waltz, F., T.-T. Nguyen, M. Arrive, A. Bochler, J. Chicher, P. Hammann, L. Kuhn, M. Quadrado, H. Mireau,Y. Hashem, and P. Giege. 2019. Small is big in Arabidopsis mitochondrial ribosome. Nat. Plants. 5:106–117. doi:10.1038/s41477-018-0339-y.

Westerfield, M. 2000. The zebrafish book. A guide for the laboratory use of zebrafish (Danio rerio). 4th ed. University of Oregon Press, Eugene.

Widenmaier, S.B., N.A. Snyder, T.B. Nguyen, A. Arduini, G.Y. Lee, A.P. Arruda, J. Saksi, A. Bartelt, and G.S. Hotamisligil. 2017. NRF1 Is an ER Membrane Sensor that Is Central to Cholesterol Homeostasis. Cell. 171:1094–1109 e15. doi:10.1016/j.cell.2017.10.003.

Yang, K., R. Huang, H. Fujihira, T. Suzuki, and N. Yan. 2018. N-glycanase NGLY1 regulates mitochondrial homeostasis and inflammation through NRF1. J Exp Med. 215:2600–2616. doi:10.1084/jem.20180783.

Yao, T., and R.E. Cohen. 2002. A cryptic protease couples deubiquitination and degradation by the proteasome. Nature. 419:403–407. doi:10.1038/nature01071.

Yeo, G., and C.B. Burge. 2004. Maximum entropy modeling of short sequence motifs with applications to RNA splicing signals. J. Comput. Biol. J. Comput. Mol. Cell Biol. 11:377–394. doi:10.1089/1066527041410418.

Yousaf, S., S.A. Sheikh, S. Riazuddin, A.M. Waryah, and Z.M. Ahmed. 2018. *INPP5K* variant causes autosomal recessive congenital cataract in a Pakistani family. Clin. Genet. 93:682–686. doi:10.1111/cge.13143.

