## Supplemental Data 1 for "Proteasome subunit *PSMC3* variants cause neurosensory syndrome combining deafness and cataract due to proteotoxic stress"

### SUPPLEMENTARY DATA

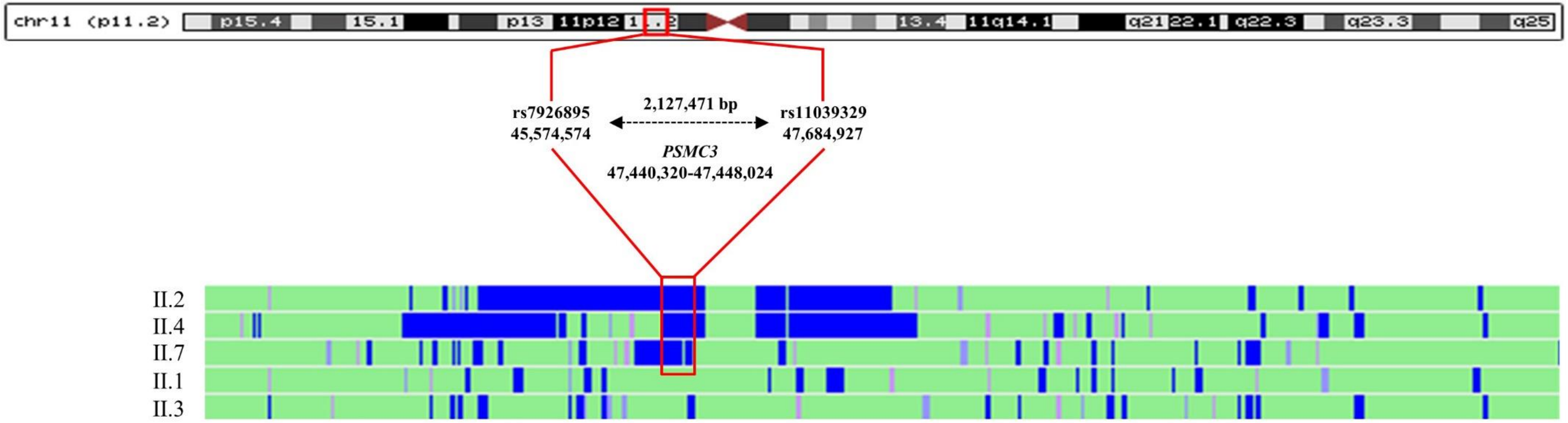


**Figure S1**. **Analysis of homozygosity region in chromosome 11**.
Homozygosity regions for three affected individuals (II.2, II.4, and II.7) and two unaffected siblings (II.1 and II.3) were analyzed using the HomoSNP software. The areas of homozygosity with >25 SNPs are colored in blue, whereas homozygosity regions defined by 15–25 consecutive SNPs are colored in pink. The 3 affected individuals share a common homozygous region between rs7926695 and rs11039329 resulting in a 2.12 Mb region encompassing the *PSMC3* gene.

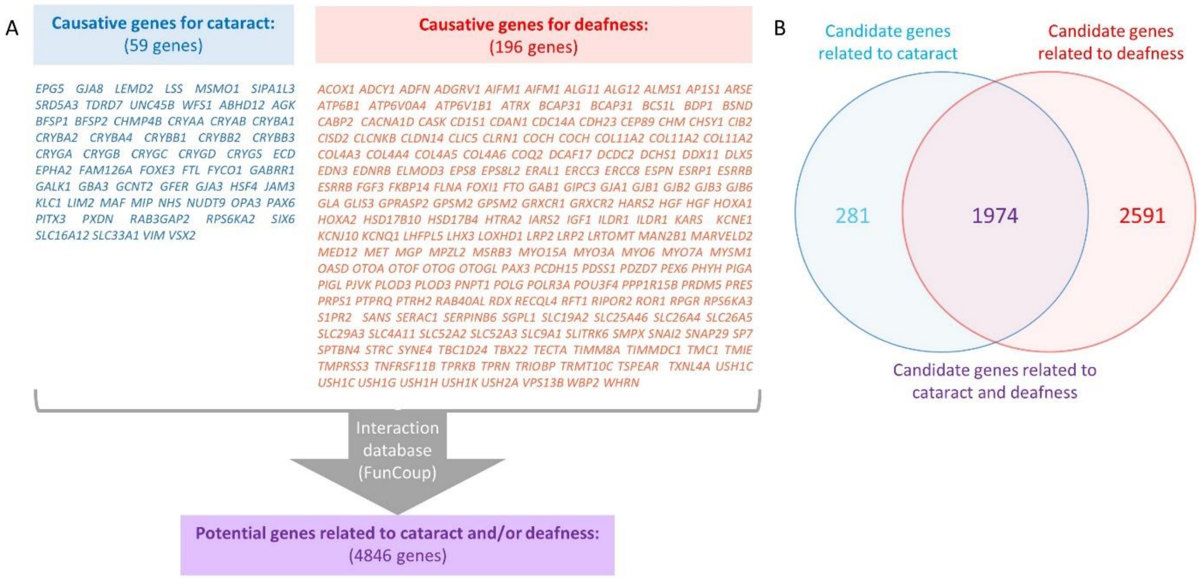


**Figure S2. Selection of candidate genes related to deafness and/or cataract.**
(A) Based on the OMIM database (Scott et al., 2018) and careful literature review two high-confidence reference gene sets including either isolated or syndromic cataract or deafness genes were defined. This resulted in 59 known cataract genes and 196 known deafness genes. Nevertheless, as the patients have congenital disorder, we excluded genes associated with ocular anomalies in which the cataract is a complication of the disease. These reference lists were used to query for potential interacting genes in the FunCoup database (4.0) (Ogris et al., 2017). Applying a confidence threshold of 0.8 and a single level of interaction a list of 4846 candidate genes related to cataract and/or deafness genes was defined.

(B) The 4846 candidate genes can be divided into 2255 candidate genes related to cataract and 4565 candidate genes related to deafness of which 1974 were in common.

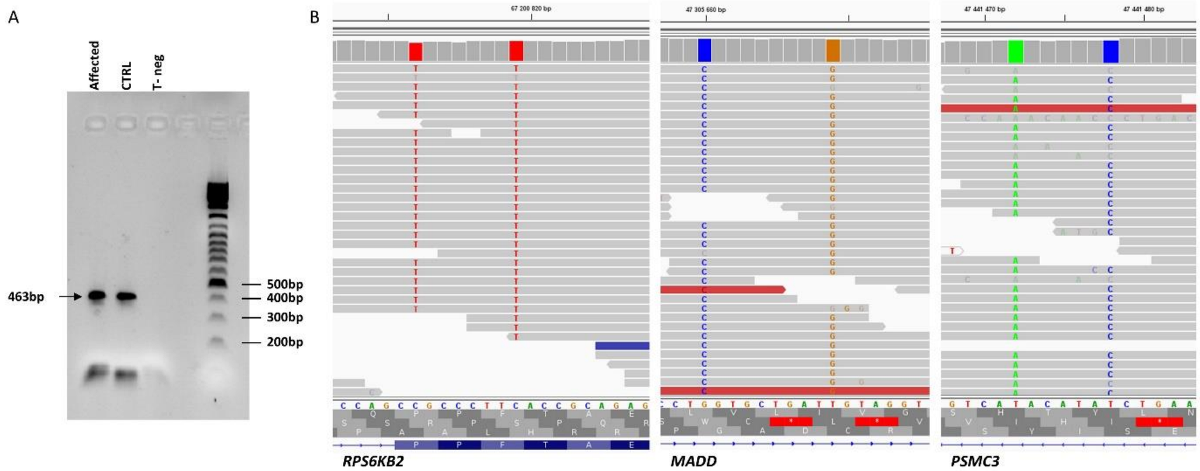


**Figure S3. Variant selection from WGS data.**

From 4,990,312 to 5,165,496 genetic variants (SNV/indel/SV) were identified per individual from the WGS analysis (Table S2). Bioinformatic analyses (see Materials and methods) highlighted 6 homozygous variations in 5 genes (*ATG13, CELF1, MADD, PSMC3* and *RPS6KB2*).

(A) Among those, the variation in *ATG13* (chr11(GRCh37):g.46680484A>G, NM_001346315.1:c.696-458A>G) is a rare (0.032% maximum allele frequency) deep intronic variant predicted to create a potential acceptor splice site. RNA analysis of *ATG13* (RTPCR on skin fibroblast from individual II.4 see Materials and Methods) surrounding intron 12 did not reveal any aberrant splicing event leaving this variant out (see below the results of the RTPCR amplification between exon 9 and exon14/15, Table S5).

(B) The *RPS6KB2* and *MADD* variations corresponded to 2 likely false positive calls. The *RPS6KB2* variant (chr11(GRCh37):g.67200812_67200819delinsTGCCCTTT) was localized at 2 highly frequent SNP: rs55987642 and rs4930427 (maximum allele frequency in gnomAD >6%). The same applies to the MADD variant (chr11(GRCh37):g.47305660_47305669delinsCGTGCTGATG) with rs12573962 and rs3816725 (maximum allele frequency in gnomAD >9%). This apply also to another variation in *PSMC3* (chr11(GRCh37):g.47441472_47441478delinsAACATAC). This can be visualized on the IGV (Thorvaldsdóttir et al., 2013). The variation in *CELF1* (Chr11(GRCh37):g.47489405T>C, NM_001330272.1:c.*4289A>G) is localized in the 3’ UTR and no prediction could be associated to it, leaving this candidate out.

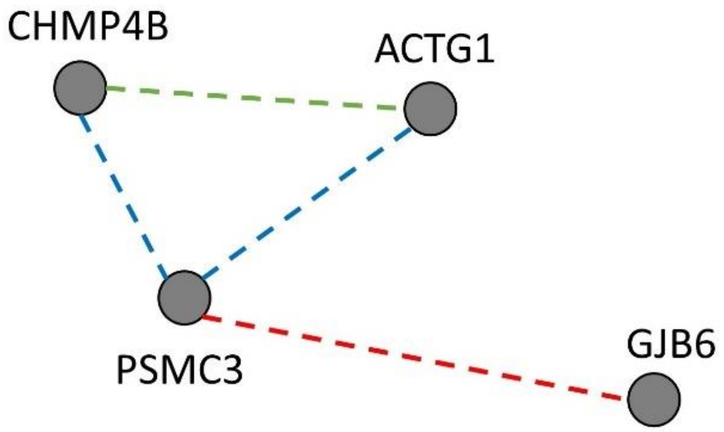


**Figure S4. Types of interactions between CHMP4B, ACTG1, GJB6 and PSMC3.**
Functional couplings displayed by FunCoup [11] are represented here with a blue line for protein-protein interaction experiment (Yeast2Hybrid), a red line for complex co-membership and a green line for a co-membership in a metabolic pathway. They are detailed in Table S4.

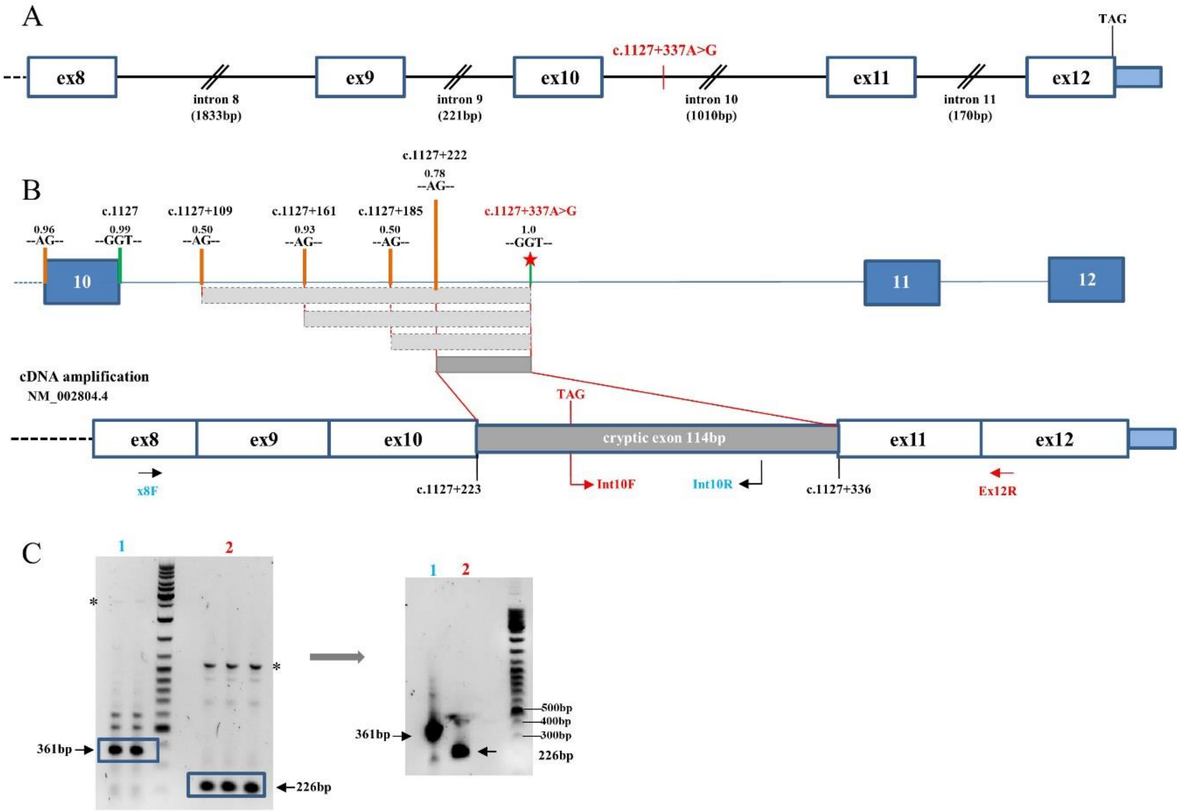


**Figure S5. Identification of the boundaries of the inserted cryptic exon.**

(A) Position of the deep intronic mutation on the *PSMC3* gene creating a strong cryptic splice donor site. Positions are given according to NM_002804.4 and positioned on the chr 11: 47,438,320-47,445,024 (11,075 bp)

(B) Schematic representation of the potential acceptor sites determined by the highest NNSPLICE score (shown below the HGVS nomenclature) as well as the resulting cryptic exons are indicated. The acceptor site at position c.1127+222 resulting in the cryptic exon displayed in dark grey was proven experimentally.

(C) Identification of the acceptor site by RT-PCR using two overlapping pairs of primers, ex8F-int10R (1) and int10F-ex12R (2). Bands of the expected sizes (361bp and 226bp, blue squares) were seen and cut for PCR reamplification (right gel) and subsequent sequencing (Figure 1). *indicates amplification of genomic DNA (bands at 2669bp and 1130bp).

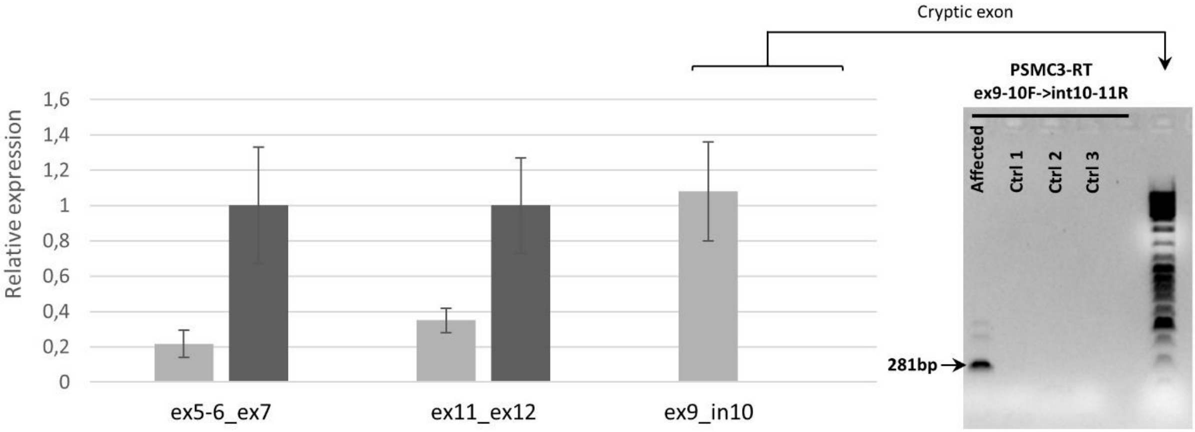


**Figure S6.** ***PSMC3*** **mRNA quantification.**

mRNA quantification between exons 5-7 and 11-12 show a significant 40/50% decrease on both fragments in the patient (II.4) as shown in light grey compared to the controls as shown in dark grey. The reference used is GAPDH. mRNA of the cryptic exon amplified using primers in exon 9 and in intron 10 was present at a high level in the patient while it was absent in the controls. The agarose electrophoresis shows the band corresponding to the cryptic exon at 281bp as amplified from the cDNA of the patient. Three independent controls of unaffected individuals were used.

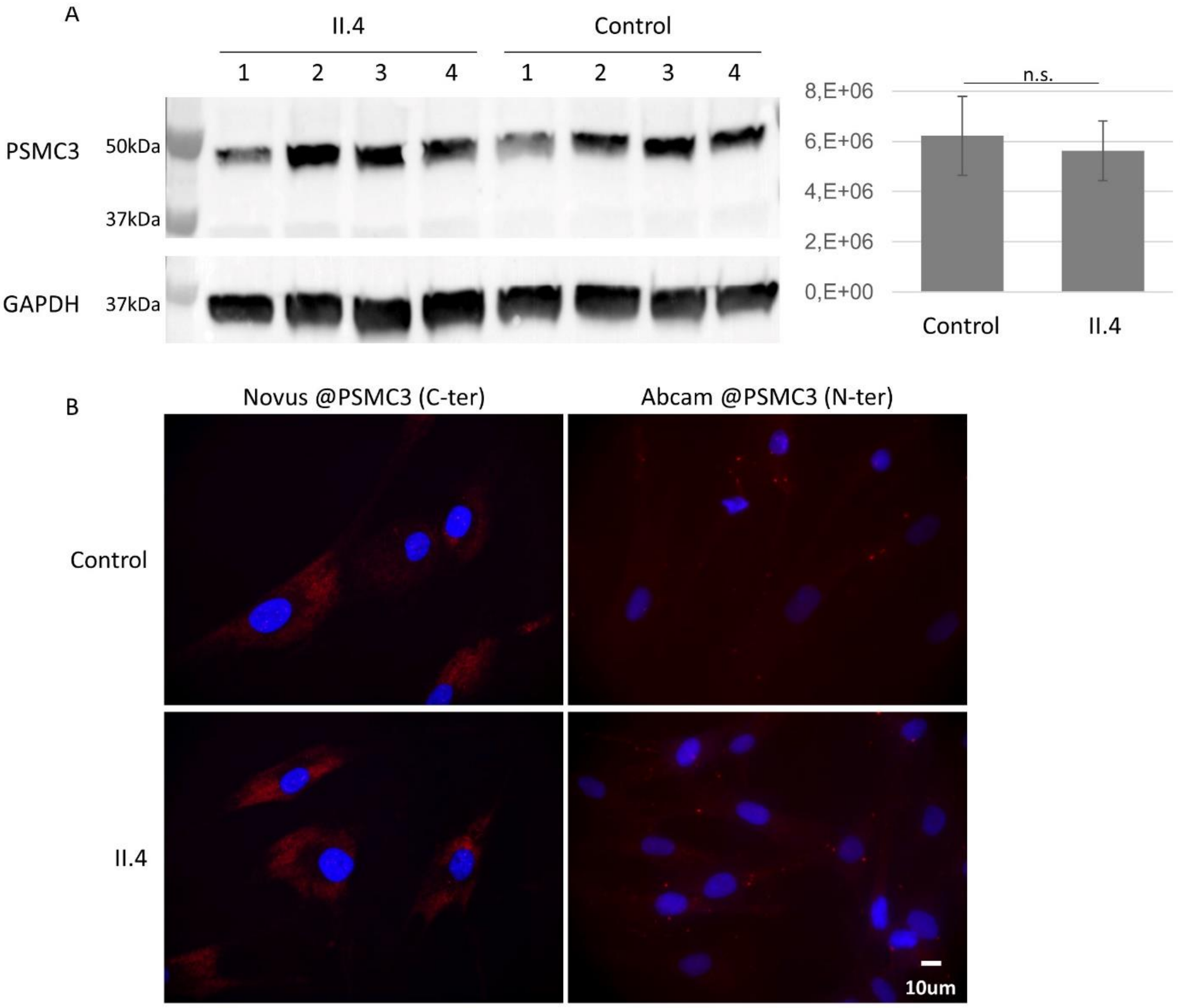


**Figure S7. No difference in PSMC3 expression or localization.**

(A) PSMC3 expression is comparable in controls and affected fibroblasts. Fibroblasts from four control individuals and from the patient II.4 (four separate batches of cells) were collected and PSMC3 detected by western blot in the whole cell lysate (Abcam antibody directed against the N-terminal thus recognizing both the full length and truncated forms of the protein). The secondary antibody was goat anti-mouse Alexa Fluor coupled 568 IgG (Invitrogen). GAPDH was used as a loading control and quantification of PSMC3 relative to GAPDH performed using Image Lab. n.s.: non-significant.

(B) No difference in PSMC3 localization: Immunofluorescence of control and affected fibroblasts (II.4) labeled with antibodies directed against the N- or C-terminal part of PSMC3 and stained with DAPI.

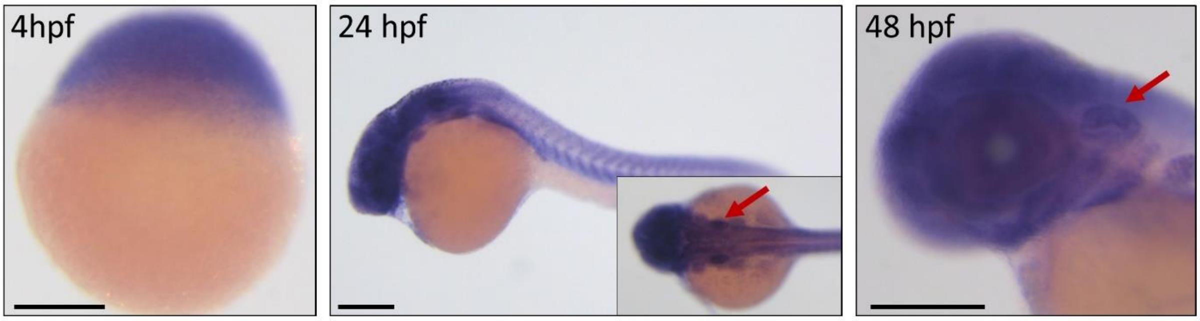


**Figure S8.** ***In situ*** **hybridization with** ***psmc3*** **antisense probe showing ubiquitous expression (4 hpf-48 hpf).** Red arrows indicate the zebrafish ear. Scale bars = 250 µm.

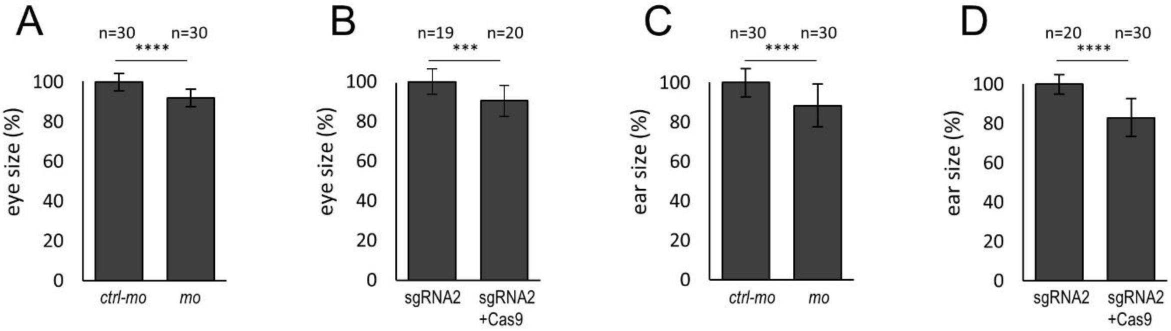


**Figure S9.** ***psmc3*** **morphants and crispants exhibit smaller lenses and inner ears.**
(A-B) Size quantification of the lens in morphants (*mo*) and F0 mutants targeted with sgRNA2 at 4 dpf. Morphants (*mo*) and crispants (sgRNA2 + Cas9) have slightly smaller lenses compared to uninjected and control injected embryos (*ctrl-mo*, sgRNA2).
(C-D) Size quantification of the inner ear in morphants (*mo*) and crispants (sgRNA2) at 4 dpf. Mo- and sgRNA2 + Cas9 injected embryos present a smaller inner ear phenotype compared to uninjected and control injected embryos (*ctrl-mo*, sgRNA2). Statistical significance was determined using the unpaired, t Test, ns = non-significant. * $p < 0.05$. ** $p < 0.01$. *** $p < 0.001$. **** $p < 0.0001$. Significance is determined relative to control injected embryos.

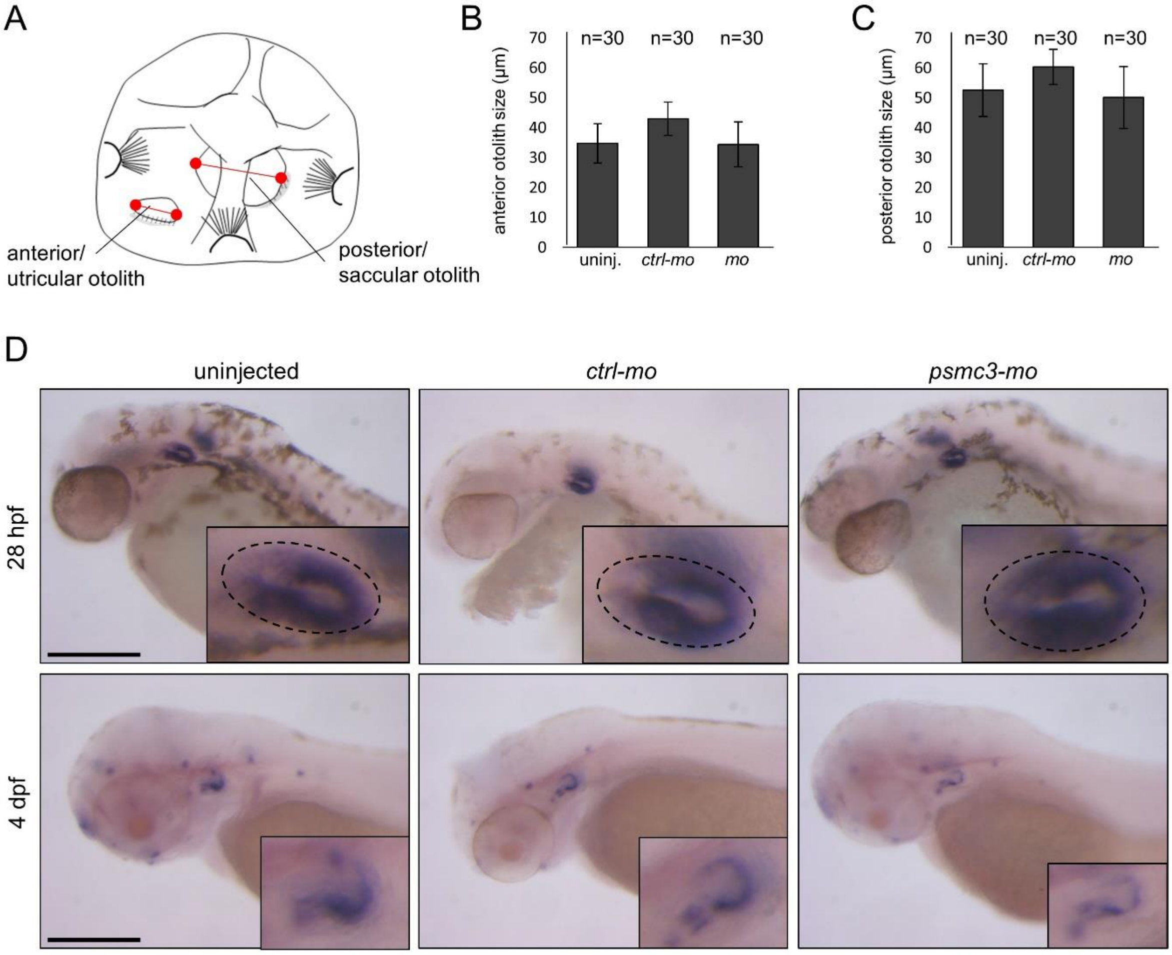


**Figure S10. Otolith development is not affected in *psmc3* morphants.**
(A) Representative image of a zebrafish ear at 4dpf. For the evaluation of otolith size diameter both otoliths were measured (longest distance, shown in red). (B-C) Otolith sizes were measured and analyzed with ImageJ. (D) Lateral views of 28 hpf and 4 dpf old embryos after morpholino injection showing expression of *otopetrin*, a gene required for otolith formation. Scale bars = 250 µm.

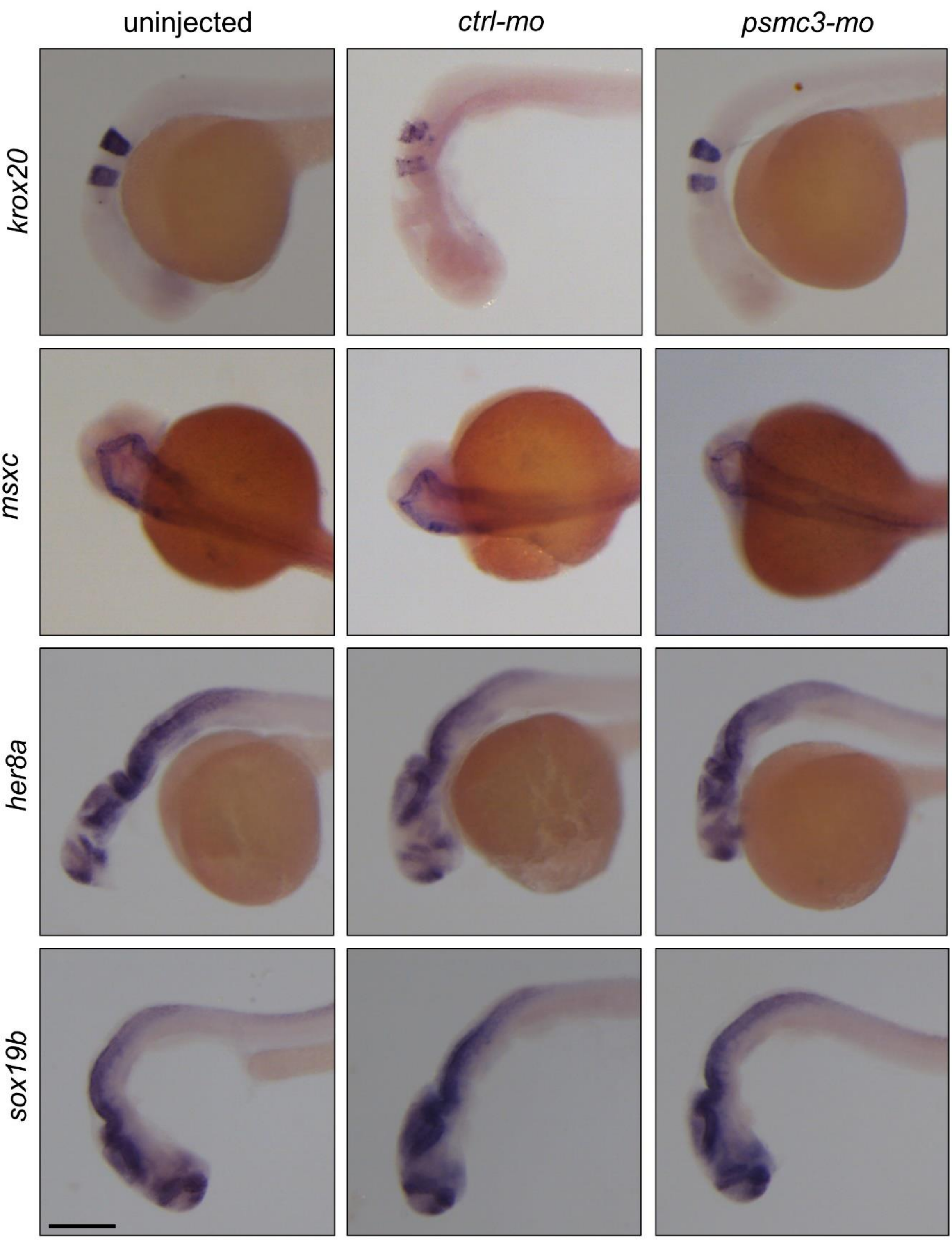


**Figure S11. *psmc3* morphants display no obvious brain malformations at 24 hpf**. Lateral views showing expression of brain markers *krox20*, *msxc*, *her8a,* and *sox19b*. Scale bar = 250 µm.

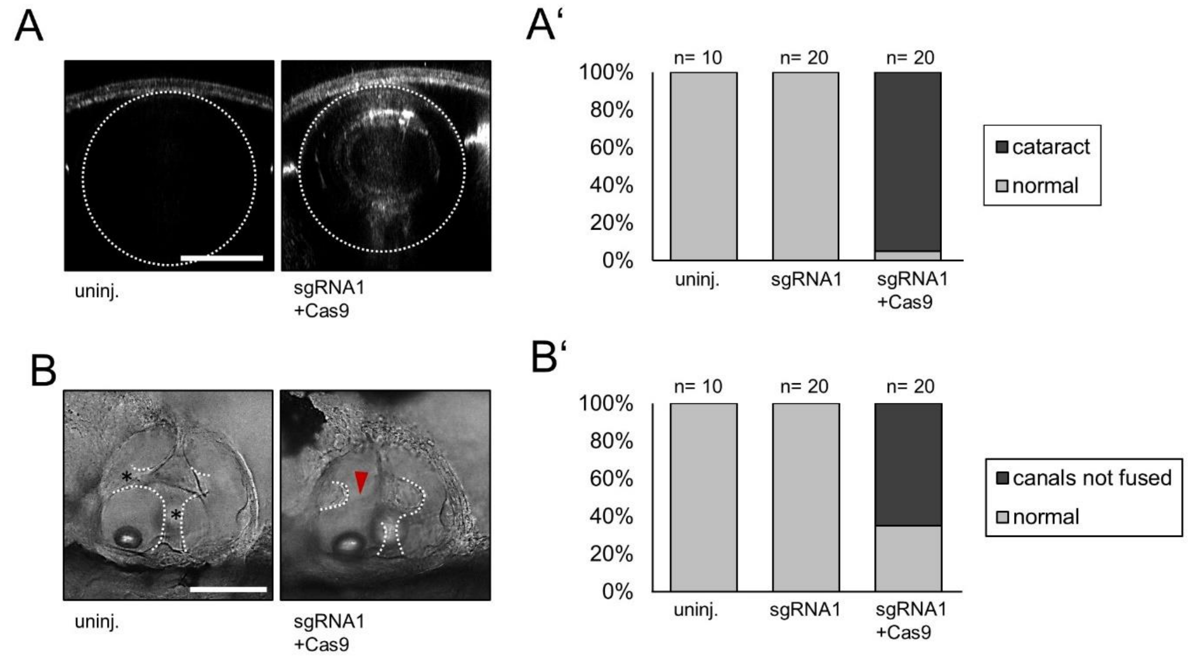


**Figure S12. A second guide RNA (sgRNA1) confirms the cataract and ear phenotype seen in morphants (*mo*) and crispants (sgRNA2).**

(A) Injection with sgRNA1 + Cas9 resulted in abnormal lens reflection but not in sgRNA injected embryos without Cas9 (sgRNA1) or uninjected embryos. Scale bar = 50 µm. (A') Quantification of embryos with abnormal lens reflection. (B) Semicircular canals were fused in 4-day-old uninjected and sgRNA1 only injected fish but not in crispants (sgRNA1+Cas9). Black asterisks indicate fused pillars. Red arrowheads mark unfused projections. Scale bar = 100 µm. (B') Quantification of embryos with an abnormal ear phenotype.

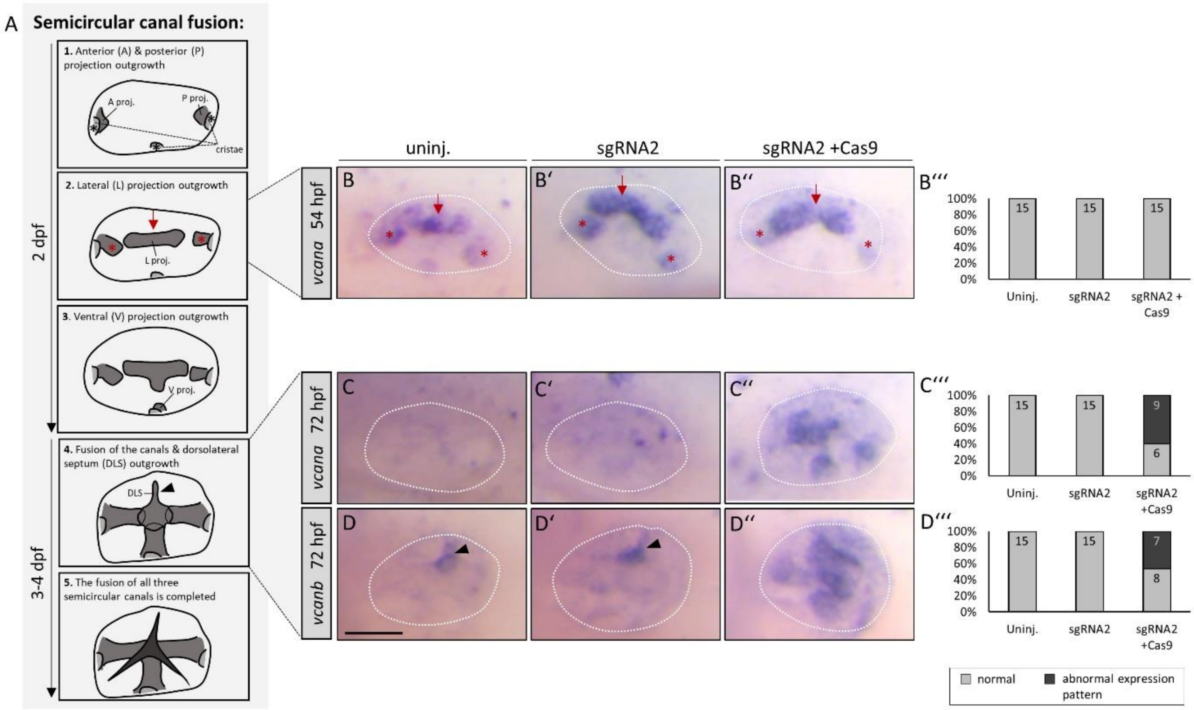


**Figure S13. Expression of genes involved in the zebrafish inner ear development in *psmc3* crispants.**

(A) Schematic representation of the semicircular canal morphogenesis between 2 and 4 dpf (inspired by Geng et al., 2013).(Geng et al., 2013) (B-B'') The expression of *versican a (vcana)* in *psmc3* crispants is normal after 54 hpf, but is (C-C'') clearly upregulated after 72 hpf compared to uninjected and control injected embryos. (D-D'') *Versican b* (*vcanb*) is expressed in the dorsolateral septum (black arrowhead), whereas in crispants it is highly upregulated in the whole unfused canal tissue. (B''', C''', D''') Quantification of embryos with an abnormal mRNA expression pattern. Scale bar = 50 µm.

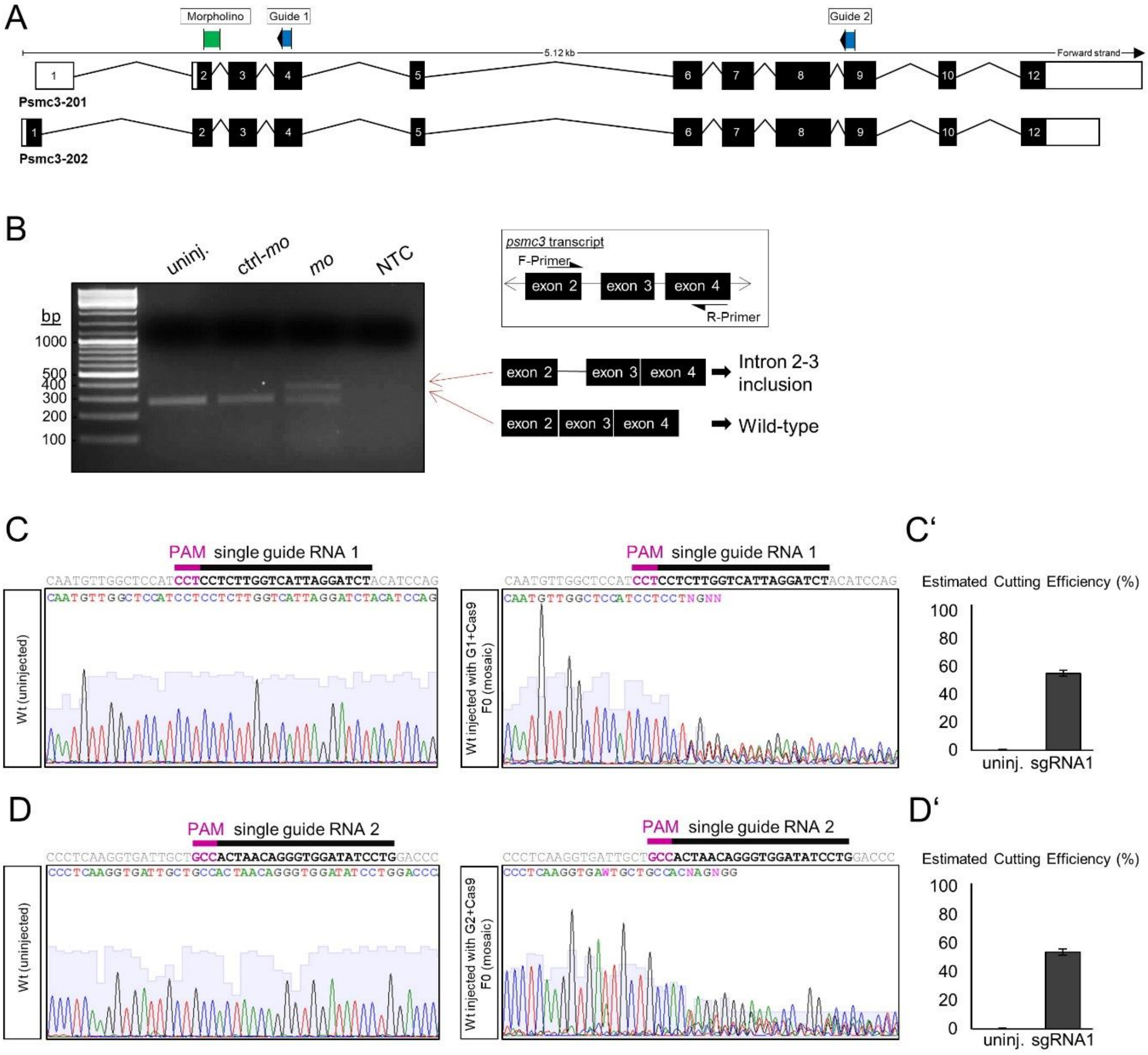


**Figure S14. Design and efficiency of morpholinos targeting *psmc3* pre-mRNA and guide RNAs targeting *psmc3*.**

**(A)** Simplified scheme of the 2 zebrafish *psmc3* isoforms. Morpholino target sequence is shown in green. Guide RNA target sequences are indicated in blue. **(B)** *psmc3-mo* injection affects splicing and leads to the inclusion of an intron. Regions amplified during PCR are shown with arrows. NTC: no template control. **(C-D)** Guide RNAs are able to cut multiple times independently during development leading to a set of small deletions and insertions. **(C)** Injection of 300 ng/µl guide RNA1 and Cas9 introduces indel in *psmc3* exon 4 with a cutting efficiency of 55.2 %. **(D)** Injection of 300 ng/µl guide RNA2 and Cas9 introduces indel in *psmc3* exon 8 with a cutting efficiency of 53.7 %. **(C'+D')** The web interface PCR-F-Seq q (http://iai-gec-server.iai.kit.edu) was used to quantify the cutting efficiency of both guide RNAs. (Etard et al., 2017)

| | II.4 | | | II.2 | | | II.7 | | | II.6 | | |
|---|---|---|---|---|---|---|---|---|---|---|---|---|
| | SNV | indel | SV | SNV | indel | SV | SNV | indel | SV | SNV | indel | SV |
| Total number of variants | 88,136 | 13,923 | 13 | 86,637 | 13,009 | 12 | 84,019 | 12,698 | 22 | 83,948 | 12,253 | 21 |
| After exclusion of variants with an allele frequency > 1% (gnomAD, 1000G, internal exome database, DGV) | 9,198 | 1,274 | 6 | 7,993 | 1,138 | 5 | 7,638 | 1,090 | 11 | 7,667 | 1,099 | 5 |
| After exclusion of SNV/indel found in the homozygous state in gnomAD and in our internal exome database | 3,138 | 507 | - | 2,991 | 433 | - | 2,843 | 414 | - | 2,834 | 424 | - |
| After exclusion of SNV/indel in 5'UTR, 3'UTR, downstream, upstream, intron and synonymous locations without local splice effect prediction | 506 | 44 | - | 525 | 34 | - | 468 | 48 | - | 473 | 44 | - |
| After exclusion of missense without SIFT, PPH2 or PhastCons prediction | 420 | 44 | - | 450 | 34 | - | 383 | 48 | - | 394 | 44 | - |
| After selection of variants consistent with recessive transmission (compound heterozygous, homozygous variants) | 1 homozygous variant (in *DGKZ*)<br>0 heterozygous compound | | | | | | | | | | | |

**Table S1. Summary of the whole exome sequencing results.**
SNV: Single nucleotide variation, indel: gain or loss of up to 50 nucleotides at a single locus, SV: Structural Variation. Annotations are gathered using Alamut Batch v1.11, especially for the variation databases including gnomAD (v2.0.2, Oct. 2017), 1000 Genomes Project phase3 release (version 20150813 v5b) and the following predictions including phastCons (UCSC, 44 vertebrates). Effect on the splice has been evaluated using the MaxEntScan (Yeo and Burge, 2004), NNSPLICE 0.9 (Reese et al., 1997) and Splice Site Finder (Shapiro and Senapathy, 1987) by calculating score change between the wild type and the mutated sequences expressed as a percent differences. Missenses have been evaluated using default parameters from PolyPhen-2 (2.2.2) (Adzhubei et al., 2010) and SIFT 4.0.3 (Kumar et al., 2009). Default cut-offs used have been described in VaRank (Geoffroy et al., 2015) for both type of predictions. Exclusion of SV with a DGV (Gold standard from 20160515) frequency > 1% is done only with studies of more than 1000 individuals.

| | II.1 | | | II.2 | | | II.3 | | | II.4 | | | II.7 | |
|---|---|---|---|---|---|---|---|---|---|---|---|---|---|---|
| | **SNV** | **indel** | **SV** | **SNV** | **indel** | **SV** | **SNV** | **indel** | **SV** | **SNV** | **indel** | **SV** | **SNV** | **indel** |
| Total number of variants | 4,013,210 | 1,147,432 | 4,376 | 4,018,208 | 1,136,593 | 10,695 | 3,966,388 | 1,097,328 | 8840 | 4,003,486 | 1,140,327 | 11,494 | 3,857,384 | 1,121,5[illegible] |
| After selection of variant found in the ROH | 3,426 | 1,250 | 6 | 2,531 | 1,101 | 10 | 3,048 | 1,148 | 6 | 2,646 | 1,065 | 9 | 2,635 | 1,1[illegible] |
| After exclusion of variants not in the cataract and deafness candidate-genes list | 816 | 304 | 3 | 652 | 312 | 4 | 723 | 292 | 2 | 666 | 287 | 2 | 663 | 2[illegible] |
| After exclusion of variants with an allele frequency >1% (gnomAD, 1000G, internal exome database, DGV) | 85 | 27 | 0 | 67 | 32 | 0 | 84 | 29 | 0 | 74 | 23 | 0 | 76 | [illegible] |
| After exclusion of SNV/indel in 5'UTR, 3'UTR, downstream, upstream, intron and synonymous locations without local splice effect prediction | 10 | 1 | - | 13 | 1 | - | 12 | 0 | - | 14 | 1 | - | 14 | [illegible] |
| After selection of homozygous variants in the affected individuals and heterozygous or absent in the healthy individuals | 6 homozygous variants in 5 genes (*ATG13, CELF1, MADD, PSMC3* and *RPS6KB2*) | | | | | | | | | | | | | |

**Table S2. Summary of the whole genome sequencing results.**

SNV: Single nucleotide variation, indel: gain or loss of up to 50 nucleotides at a single locus, SV: Structural Variation. ROH: Region of homozygosity as defined by the SNParrays. Annotations are gathered using Alamut Batch v1.11, especially for the variation databases including gnomAD (v2.0.2, Oct. 2017), 1000 Genomes Project phase3 release (version 20150813 v5b) and the following predictions including phastCons (UCSC, 44 vertebrates). Effect on the splice has been evaluated using the MaxEntScan (Yeo and Burge, 2004), NNSPLICE 0.9 (Reese et al., 1997) and Splice Site Finder (Shapiro and Senapathy, 1987) by calculating score change between the wild type and the mutated sequences expressed as a percent differences. Missenses have been evaluated using default parameters from PolyPhen-2 (2.2.2) (Adzhubei et al., 2010) and SIFT 4.0.3 (Kumar et al., 2009). Default cut-offs used have been described in VaRank (Geoffroy et al., 2015) for both type of predictions. Exclusion of SV with a DGV (Gold standard from 20160515) frequency > 1% is done only with studies of more than 1000 individuals.

| **HGNC approved name** | **Human** | **Mouse** | **Yeast** |
|---|---|---|---|
| Proteasome 26S subunit, ATPase 3 | PSMC3 (PRS6A_HUMAN) | psmc3 (PRS6A_MOUSE*) | RPT5 (YOR117W) |
| Actin gamma 1 | ACTG1 (ACTG_HUMAN) | actg1 (ACTG_MOUSE) | ACT1 (YFL039C) |
| Charged multivesicular body protein 4B | CHMP4B (CHM4B_HUMAN) | chmp4b (CHM4B_MOUSE) | SNF7 (YLR025W) |
| Gap junction protein beta 6 | GJB6 (CXB6_HUMAN) | gjb6 (CXB6_MOUSE) | |

**Table S3. Orthologous ID equivalent of *PSMC3*, *ACTG1*, *CHMP4B* and *GJB6* in human, mouse and yeast.**
For each species, official gene symbols are indicated with the Uniprot (The UniProt Consortium, 2016) entry name into parenthesis. HGNC: Hugo Gene Nomenclature Committee. * To avoid confusion, we would like to highlight that in 2006 Binato *et al* (Binato et al., 2006) named the mouse orthologue of *PSMC3* “PRSA_MOUSE” while it is currently named “PRS6A_MOUSE”.

| Human protein interaction | Publication showing the interaction | Organism of interest | Orthologous genes names used |
|---|---|---|---|
| PSMC3 / GJB6 | Binato et al. (Binato et al., 2006) | Mouse | PRSA_MOUSE / CXB6_MOUSE |
| PSMC3 / ACTG1 | Guerrero et al. (Guerrero et al., 2008) | Yeast | *RPT5 / ACT1* |
| PSMC3 / CHMP4B | Wang et al. (Wang et al., 2012) | Yeast | YOR117W.1 / YLR025W.2 |

**Table S4. Publications showing genes interactions between *PSMC3*, *ACTG1*, *CHMP4B* and *GJB6*.**

| | | | RAW Spectral Count | | | | | | | | | NORMALIZED Spectral Count | | | | | | Mean #spectra | | STDEV #spectra | | Ratio (Mean) |
|---|---|---|---|---|---|---|---|---|---|---|---|---|---|---|---|---|---|---|---|---|---|---|
| | | | CONTROL | | | HEALTHY | | | AFFECTED | | | HEALTHY | | | AFFECTED | | | | | | | |
| UniProt Acc | Gene name | Description from UniProt | #1 | #2 | #3 | #1 | #2 | #3 | #1 | #2 | #3 | #1 | #2 | #3 | #1 | #2 | #3 | H | A | H | A | A/H |
| PSA1_HUMAN | *PSMA1* | Proteasome subunit alpha type-1 | 0 | 0 | 0 | 41 | 44 | 49 | 64 | 61 | 57 | 44.02 | 43.29 | 48.06 | 63.49 | 60.78 | 55.83 | 45.1 | 60.0 | 2.6 | 3.9 | 1.33 |
| PSA2_HUMAN | *PSMA2* | Proteasome subunit alpha type-2 | 0 | 0 | 0 | 27 | 28 | 25 | 33 | 31 | 32 | 28.99 | 27.55 | 24.52 | 32.73 | 30.89 | 31.35 | 27.0 | 31.7 | 2.3 | 1.0 | 1.17 |
| PSA3_HUMAN | *PSMA3* | Proteasome subunit alpha type-3 | 0 | 0 | 0 | 28 | 27 | 27 | 43 | 41 | 44 | 30.06 | 26.56 | 26.48 | 42.65 | 40.85 | 43.10 | 27.7 | 42.2 | 2.0 | 1.2 | 1.52 |
| PSA4_HUMAN | *PSMA4* | Proteasome subunit alpha type-4 | 0 | 0 | 0 | 31 | 31 | 35 | 51 | 47 | 46 | 33.28 | 30.50 | 34.33 | 50.59 | 46.83 | 45.06 | 32.7 | 47.5 | 2.0 | 2.8 | 1.45 |
| PSA5_HUMAN | *PSMA5* | Proteasome subunit alpha type-5 | 0 | 0 | 0 | 14 | 18 | 16 | 24 | 20 | 20 | 15.03 | 17.71 | 15.69 | 23.81 | 19.93 | 19.59 | 16.1 | 21.1 | 1.4 | 2.3 | 1.31 |
| PSA6_HUMAN | *PSMA6* | Proteasome subunit alpha type-6 | 0 | 0 | 0 | 32 | 37 | 43 | 49 | 56 | 56 | 34.36 | 36.40 | 42.17 | 48.61 | 55.80 | 54.85 | 37.6 | 53.1 | 4.1 | 3.9 | 1.41 |
| PSA7_HUMAN | *PSMA7* | Proteasome subunit alpha type-7 | 0 | 0 | 0 | 33 | 40 | 42 | 60 | 54 | 49 | 35.43 | 39.36 | 41.19 | 59.52 | 53.81 | 48.00 | 38.7 | 53.8 | 2.9 | 5.8 | 1.39 |
| PSB1_HUMAN | *PSMB1* | Proteasome subunit beta type-1 | 0 | 0 | 0 | 26 | 31 | 35 | 41 | 45 | 43 | 27.92 | 30.50 | 34.33 | 40.67 | 44.84 | 42.12 | 30.9 | 42.5 | 3.2 | 2.1 | 1.38 |
| PSB2_HUMAN | *PSMB2* | Proteasome subunit beta type-2 | 0 | 0 | 0 | 13 | 14 | 15 | 31 | 31 | 25 | 13.96 | 13.77 | 14.71 | 30.75 | 30.89 | 24.49 | 14.1 | 28.7 | 0.5 | 3.7 | 2.03 |
| PSB3_HUMAN | *PSMB3* | Proteasome subunit beta type-3 | 0 | 0 | 0 | 15 | 21 | 18 | 29 | 26 | 25 | 16.10 | 20.66 | 17.65 | 28.77 | 25.91 | 24.49 | 18.1 | 26.4 | 2.3 | 2.2 | 1.45 |
| PSB4_HUMAN | *PSMB4* | Proteasome subunit beta type-4 | 0 | 0 | 0 | 13 | 12 | 12 | 30 | 28 | 24 | 13.96 | 11.81 | 11.77 | 29.76 | 27.90 | 23.51 | 12.5 | 27.1 | 1.3 | 3.2 | 2.16 |
| PSB5_HUMAN | *PSMB5* | Proteasome subunit beta type-5 | 0 | 1 | 0 | 25 | 27 | 27 | 39 | 35 | 35 | 26.84 | 26.56 | 26.48 | 38.69 | 34.87 | 34.28 | 26.6 | 35.9 | 0.2 | 2.4 | 1.35 |
| PSB6_HUMAN | *PSMB6* | Proteasome subunit beta type-6 | 0 | 0 | 0 | 6 | 8 | 9 | 17 | 16 | 11 | 6.44 | 7.87 | 8.83 | 16.86 | 15.94 | 10.77 | 7.7 | 14.5 | 1.2 | 3.3 | 1.88 |
| PSB7_HUMAN | *PSMB7* | Proteasome subunit beta type-7 | 0 | 0 | 0 | 8 | 9 | 9 | 11 | 14 | 11 | 8.59 | 8.85 | 8.83 | 10.91 | 13.95 | 10.77 | 8.8 | 11.9 | 0.1 | 1.8 | 1.36 |
| PSB8_HUMAN | *PSMB8* | Proteasome subunit beta type-8 | 0 | 0 | 0 | 10 | 12 | 14 | 21 | 19 | 15 | 10.74 | 11.81 | 13.73 | 20.83 | 18.93 | 14.69 | 12.1 | 18.2 | 1.5 | 3.1 | 1.50 |
| PSB9_HUMAN | *PSMB9* | Proteasome subunit beta type-9 | 0 | 0 | 0 | 6 | 5 | 5 | 6 | 5 | 7 | 6.44 | 4.92 | 4.90 | 5.95 | 4.98 | 6.86 | 5.4 | 5.9 | 0.9 | 0.9 | 1.09 |
| PSB10_HUMAN | *PSMB10* | Proteasome subunit beta type-10 | 0 | 0 | 0 | 4 | 6 | 6 | 4 | 5 | 6 | 4.29 | 5.90 | 5.88 | 3.97 | 4.98 | 5.88 | 5.4 | 4.9 | 0.9 | 1.0 | 0.92 |
| PRS4_HUMAN | *PSMC1* | 26S proteasome regulatory subunit 4 | 0 | 0 | 0 | 63 | 72 | 71 | 84 | 77 | 74 | 67.64 | 70.84 | 69.63 | 83.32 | 76.72 | 72.49 | 69.4 | 77.5 | 1.6 | 5.5 | 1.12 |
| PRS7_HUMAN | *PSMC2* | 26S proteasome regulatory subunit 7 | 0 | 0 | 0 | 76 | 94 | 84 | 112 | 108 | 102 | 81.60 | 92.49 | 82.38 | 111.10 | 107.61 | 99.91 | 85.5 | 106.2 | 6.1 | 5.7 | 1.24 |
| PRS6A_HUMAN | *PSMC3* | 26S proteasome regulatory subunit 6A | 0 | 0 | 0 | 97 | 108 | 107 | 104 | 100 | 103 | 104.15 | 106.26 | 104.94 | 103.16 | 99.64 | 100.89 | 105.1 | 101.2 | 1.1 | 1.8 | 0.96 |
| PRS6B_HUMAN | *PSMC4* | 26S proteasome regulatory subunit 6B | 0 | 0 | 0 | 85 | 86 | 99 | 103 | 96 | 104 | 91.26 | 84.61 | 97.09 | 102.17 | 95.66 | 101.87 | 91.0 | 99.9 | 6.2 | 3.7 | 1.10 |
| PRS8_HUMAN | *PSMC5* | 26S proteasome regulatory subunit 8 | 0 | 0 | 0 | 68 | 65 | 72 | 92 | 83 | 84 | 73.01 | 63.95 | 70.61 | 91.26 | 82.70 | 82.28 | 69.2 | 85.4 | 4.7 | 5.1 | 1.23 |
| PRS10_HUMAN | *PSMC6* | 26S proteasome regulatory subunit 10B | 0 | 0 | 0 | 89 | 88 | 92 | 88 | 87 | 96 | 95.56 | 86.58 | 90.23 | 87.29 | 86.69 | 94.04 | 90.8 | 89.3 | 4.5 | 4.1 | 0.98 |
| PSMD1_HUMAN | *PSMD1* | 26S proteasome non-ATPase regulatory subunit 1 | 0 | 0 | 0 | 97 | 109 | 104 | 133 | 129 | 113 | 104.15 | 107.24 | 102.00 | 131.93 | 128.54 | 110.69 | 104.5 | 123.7 | 2.6 | 11.4 | 1.18 |
| PSMD2_HUMAN | *PSMD2* | 26S proteasome non-ATPase regulatory subunit 2 | 0 | 0 | 0 | 105 | 124 | 119 | 127 | 136 | 133 | 112.73 | 122.00 | 116.71 | 125.98 | 135.51 | 130.28 | 117.1 | 130.6 | 4.6 | 4.8 | 1.11 |
| PSMD3_HUMAN | *PSMD3* | 26S proteasome non-ATPase regulatory subunit 3 | 0 | 0 | 0 | 74 | 83 | 81 | 109 | 103 | 97 | 79.45 | 81.66 | 79.44 | 108.12 | 102.63 | 95.02 | 80.2 | 101.9 | 1.3 | 6.6 | 1.27 |
| PSMD4_HUMAN | *PSMD4* | 26S proteasome non-ATPase regulatory subunit 4 | 0 | 0 | 0 | 27 | 25 | 27 | 38 | 32 | 30 | 28.99 | 24.60 | 26.48 | 37.69 | 31.89 | 29.39 | 26.7 | 33.0 | 2.2 | 4.3 | 1.24 |
| PSMD5_HUMAN | *PSMD5* | 26S proteasome non-ATPase regulatory subunit 5 | 0 | 0 | 0 | 21 | 22 | 19 | 20 | 20 | 15 | 22.55 | 21.65 | 18.63 | 19.84 | 19.93 | 14.69 | 20.9 | 18.2 | 2.0 | 3.0 | 0.87 |
| PSMD6_HUMAN | *PSMD6* | 26S proteasome non-ATPase regulatory subunit 6 | 0 | 0 | 0 | 54 | 68 | 72 | 71 | 73 | 68 | 57.98 | 66.90 | 70.61 | 70.43 | 72.74 | 66.61 | 65.2 | 69.9 | 6.5 | 3.1 | 1.07 |
| PSMD7_HUMAN | *PSMD7* | 26S proteasome non-ATPase regulatory subunit 7 | 0 | 0 | 0 | 34 | 46 | 36 | 42 | 40 | 35 | 36.50 | 45.26 | 35.31 | 41.66 | 39.86 | 34.28 | 39.0 | 38.6 | 5.4 | 3.8 | 0.99 |
| PSMD8_HUMAN | *PSMD8* | 26S proteasome non-ATPase regulatory subunit 8 | 0 | 0 | 0 | 19 | 24 | 25 | 24 | 23 | 21 | 20.40 | 23.61 | 24.52 | 23.81 | 22.92 | 20.57 | 22.8 | 22.4 | 2.2 | 1.7 | 0.98 |
| PSMD9_HUMAN | *PSMD9* | 26S proteasome non-ATPase regulatory subunit 9 | 0 | 0 | 0 | 27 | 33 | 32 | 27 | 24 | 22 | 28.99 | 32.47 | 31.38 | 26.78 | 23.91 | 21.55 | 30.9 | 24.1 | 1.8 | 2.6 | 0.78 |
| PSD10_HUMAN | *PSMD10* | 26S proteasome non-ATPase regulatory subunit 10 | 0 | 0 | 0 | 4 | 5 | 5 | 4 | 2 | 2 | 4.29 | 4.92 | 4.90 | 3.97 | 1.99 | 1.96 | 4.7 | 2.6 | 0.4 | 1.2 | 0.56 |
| PSD11_HUMAN | *PSMD11* | 26S proteasome non-ATPase regulatory subunit 11 | 0 | 0 | 0 | 73 | 79 | 81 | 93 | 95 | 88 | 78.38 | 77.73 | 79.44 | 92.25 | 94.66 | 86.20 | 78.5 | 91.0 | 0.9 | 4.4 | 1.16 |
| PSD12_HUMAN | *PSMD12* | 26S proteasome non-ATPase regulatory subunit 12 | 0 | 0 | 0 | 54 | 62 | 62 | 62 | 66 | 57 | 57.98 | 61.00 | 60.81 | 61.50 | 65.76 | 55.83 | 59.9 | 61.0 | 1.7 | 5.0 | 1.02 |
| PSD13_HUMAN | *PSMD13* | 26S proteasome non-ATPase regulatory subunit 13 | 0 | 0 | 0 | 45 | 53 | 53 | 63 | 60 | 48 | 48.31 | 52.15 | 51.98 | 62.49 | 59.79 | 47.02 | 50.8 | 56.4 | 2.2 | 8.3 | 1.11 |
| PSDE_HUMAN | *PSMD14* | 26S proteasome non-ATPase regulatory subunit 14 | 0 | 0 | 0 | 35 | 35 | 36 | 34 | 34 | 32 | 37.58 | 34.44 | 35.31 | 33.73 | 33.88 | 31.35 | 35.8 | 33.0 | 1.6 | 1.4 | 0.92 |

**Table S5. Proteasome proteins identified using nanoLC-MS/MS analysis and quantified by Spectral Count.**

PSMC3 was immunoprecipitated from control (CONTROL#1/2/3) and patients (HEALTHY#1/2/3 and AFFECTED#1/2/3) fibroblast protein lysates. Interaction partners were determined by searching against the complete UniProtKB Human proteome set (The UniProt Consortium, 2016). The total number of MS/MS fragmentation spectra identified by Mascot algorithm (Perkins et al., 1999) and validated at FDR<1% were then counted for each protein in each sample. Spectral Count values were then normalized by the total number of spectra in each sample. in order to calculate the final relative abundance of each protein between the AFFECTED triplicates and the HEALTHY triplicates. H: Healthy, A: Affected.

| Organism | Primer name | Sequence (5' -> 3') | cDNA position |
|---|---|---|---|
| Human | ATG13-RT-ex9F | CAGGAAACAAGGGCATGAAT | c.423 to c.442 |
| | ATG13-RT-ex14R-15R | AGGCGAGAGCTTGAGAG*TTG* | c.886 to c.905 |
| | PSMC3-ex10F | CTCAGGTAATTGCAGCCACA | c.982-5 to c.996 |
| | PSMC3-int10R | CGCCTGTAGTCCCAGCTACT | c.1127+514 to c.1127+533 |
| | PSMC3-int10R | GCGACAGAGCGAGACACTG | c.1127+410 to c.1127+428 |
| | PSMC3-RT-ex9-10F | AAGTTAAG*GTAATTGCAGCCACA* | c.974 to c.996 |
| | PSMC3-RT-ex12R | CATGTAGTCCTCGTGGGTGA | c.1244 to c.1263 |
| | PSMC3-RT-int10F | ACAGAGGCTGGAGGCACTTA | intron 10 c.1127+245 to c.1127+264 |
| | PSMC3-RT-int10R1 | TTCAGATGAGGGGTGGAGTC | intron 10 c.1127+222 to c.1127+241 |
| | PSMC3-RT-ex11R | GCCAGCTCCTCGTAGTTCAC | c.1135 to c.1154 |
| | PSMC3-RT-int10-ex11R | TCACGTCAGGA*CTGAACTCAA* | c.1138>c.1128 to c.1127+336-c.1127+327 |
| | PSMC3-RT-QPCR-ex5-6F | TAAAGCCAGGAGACCTGGTG*GGTG* | c.434 to c.457 |
| | PSMC3-RT-QPCR-ex7R | GCCCATACATCAGCACCCCTTT | c.661 to c.682 |
| | PSMC3-RT-QPCR-ex11F | GACGTGAACTACGAGGAGCTG | c.1132 to c.1152 |
| | PSMC3-RT-QPCR-ex12Rbis | GCACTTCAGCCGTGAGACT | c*19 to c*37 |
| Zebrafish | psmc3-inSitu-ex2F | GAAGAGATTGTTCAGAGGACTCG | c.79>c.101 |
| | psmc3-inSitu-ex9R | GGGCATAGGGAACTCAATCTTA | c.1014>c.1035 |
| | psmc3-full_length-ex1F | GCAGAATTCATGTCGTCGCTGAATGACAGA | c.1>c.21 |
| | psmc3-full_length-ex11R | TTAGCGGCCGCTTAAGCATAGTACTGCAAATT | c.1270>c.1284 |
| | psmc3-mo-RT-ex2F | GAAGAGATTGTTCAGAGGACTCG | c.79>c.101 |
| | psmc3-mo-RT-ex4R | TGCCGTGTGGAGGTTTTGAT | c.334>c.353 |
| | psmc3-guide1-F | GGCTGGAGTATGGTGTTAAAGC | intron 3 c.249+46>c.249+67 |
| | psmc3-guide1-R | AAAGATGGATGGAAGAATTTGG | intron 4 c.345+38>c.345+59 |
| | psmc3-guide2-ex8F | TGGAGCTGCTCAATCAGTTAGA | c.895>c.917 |
| | psmc3-guide2-R | GGGCATAGGGAACTCAATCTTA | c.1013>c.1035 |

**Table S6. Primers used in this study.** All positions and sequence are given for human according to the RefSeq Gene (O'Leary et al., 2016) identifiers including NM_002804.4 for *PSMC3* and NM_001346315.1 for *ATG13* or for zebrafish in the Zfin database (Ruzicka et al., 2018) (ZDB-GENE-030131-666) or ensembl database (Zerbino et al., 2017) (ENSDART00000171691.2).
For primers that are overlapping 2 exons, the sequence of the following exon is in italic.

| **Primary antibody name** | **Applications** | **Producer** | **Conditions** |
|---|---|---|---|
| Rabbit anti-PSMC3 | WB. IF | Novus Biologicals NBP1-86962 | WB: 1/1000 5% non-fat dry milk. IF: 1/500 |
| Rabbit anti-PSMC3 | WB. IF. IP. MS | Abcam ab171969 | WB: 1/2000. 5% non-fat dry milk. IP/MS 10µl/mg proteins; IF: 1/100 |
| Rabbit anti-ACTG1 | WB | Novus Biologicals NB600-533SS | WB: 1/10 000. 5% non-fat dry milk |
| Rabbit anti-GAPDH | WB | Abcam #ab181602 | WB: 1/5000. 5% non-fat dry milk |
| Mouse anti-ubiquitin | WB | Invitrogen #13-1600 | WB: 1/1000 |
| Mouse Anti-acetylated Tubulin | IF | Sigma-Aldrich T7451 | IF:1/500 |
| **Secondary antibody name** | | | |
| Goat anti-Rabbit Alexa Fluor 568 | IF | ThermoFisher Scientific A-11011 | IF: 1/750 |
| Donkey anti-Rabbit HRP ECL Rabbit IgG. HRP-linked | WB | GE Healthcare Life Sciences NA934V | WB: 1/5000 5% non-fat dry milk |
| Mouse anti-Rabbit | WB | Cell Signalling #5127 | WB: 1/2000 5% non-fat dry milk |

**Table S7. List of antibodies used in this study.** IF: immunofluorescence. IP: immunoprecipitation. MS: mass spectrometry. WB: Western Blot.

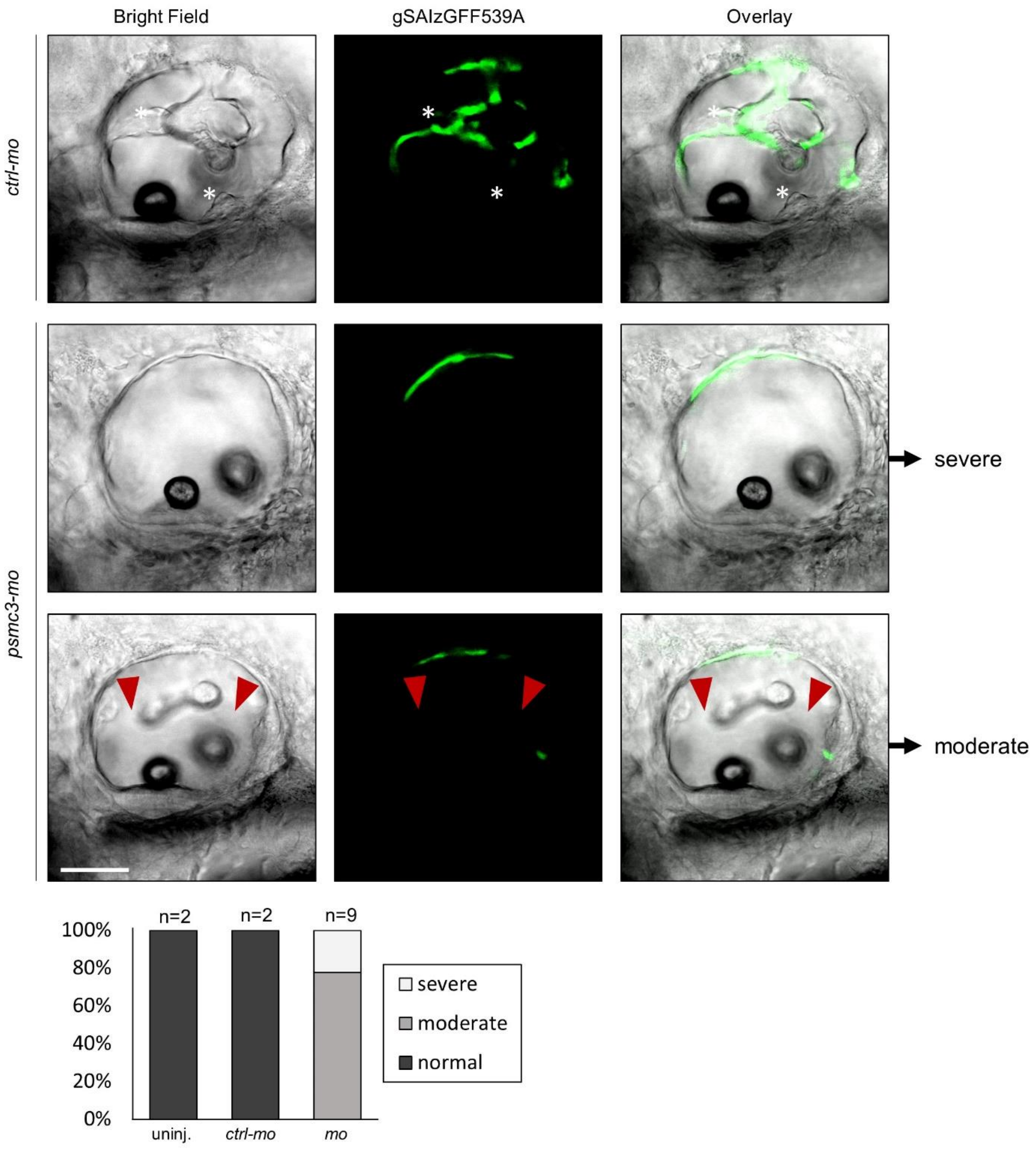


**Movie S1. Life imaging of semicircular canal formation (56 and 72 hpf) in *ctrl-mo* and *mo* injected embryos.**

Morpholino (MO)-mediated knock-down of Psmc3 resulted in abnormal ear development. Canal pillars form between 56-72 hpf in uninjected and control injected fish (*ctrl-mo*) but not in morphants (*mo*). White asterisks indicate fused pillars. Red arrowheads mark unfused projections. Severe = no outgrowth of projections of semicircular canals. Moderate = projections do not fuse. Scale bar = 50 µm.

### SUPPLEMENTARY REFERENCES


Adzhubei, I.A., S. Schmidt, L. Peshkin, V.E. Ramensky, A. Gerasimova, P. Bork, A.S. Kondrashov, and S.R. Sunyaev. 2010. A method and server for predicting damaging missense mutations. *Nat. Methods*. 7:248–249. doi:10.1038/nmeth0410-248.

Binato, R., C.E. Alvarez Martinez, L. Pizzatti, B. Robert, and E. Abdelhay. 2006. SMAD 8 binding to mice <em>Msx1</em> basal promoter is required for transcriptional activation. *Biochem. J.* 393:141. doi:10.1042/BJ20050327.

Etard, C., S. Joshi, J. Stegmaier, R. Mikut, and U. Strähle. 2017. Tracking of Indels by DEcomposition is a Simple and Effective Method to Assess Efficiency of Guide RNAs in Zebrafish. *Zebrafish*. 14:586–588. doi:10.1089/zeb.2017.1454.

Geng, F.-S., L. Abbas, S. Baxendale, C.J. Holdsworth, A.G. Swanson, K. Slanchev, M. Hammerschmidt, J. Topczewski, and T.T. Whitfield. 2013. Semicircular canal morphogenesis in the zebrafish inner ear requires the function of gpr126 (lauscher), an adhesion class G protein-coupled receptor gene. *Development*. 140:4362–4374. doi:10.1242/dev.098061.

Geoffroy, V., C. Pizot, C. Redin, A. Piton, N. Vasli, C. Stoetzel, A. Blavier, J. Laporte, and J. Muller. 2015. VaRank: a simple and powerful tool for ranking genetic variants. *PeerJ*. 3:e796. doi:10.7717/peerj.796.

Guerrero, C., T. Milenković, N. Pržulj, P. Kaiser, and L. Huang. 2008. Characterization of the proteasome interaction network using a QTAX-based tag-team strategy and protein interaction network analysis. *Proc. Natl. Acad. Sci.* 105:13333. doi:10.1073/pnas.0801870105.

Kumar, P., S. Henikoff, and P.C. Ng. 2009. Predicting the effects of coding non-synonymous variants on protein function using the SIFT algorithm. *Nat. Protoc.* 4:1073–1081. doi:10.1038/nprot.2009.86.

Ogris, C., D. Guala, M. Kaduk, and E.L.L. Sonnhammer. 2017. FunCoup 4: new species, data, and visualization. *Nucleic Acids Res.* 46:D601–D607. doi:10.1093/nar/gkx1138.

O'Leary, N.A., M.W. Wright, J.R. Brister, S. Ciufo, D. Haddad, R. McVeigh, B. Rajput, B. Robbertse, B. Smith-White, D. Ako-Adjei, A. Astashyn, A. Badretdin, Y. Bao, O. Blinkova, V. Brover, V. Chetvernin, J. Choi, E. Cox, O. Ermolaeva, C.M. Farrell, T. Goldfarb, T. Gupta, D. Haft, E. Hatcher, W. Hlavina, V.S. Joardar, V.K. Kodali, W. Li, D. Maglott, P. Masterson, K.M. McGarvey, M.R. Murphy, K. O'Neill, S. Pujar, S.H. Rangwala, D. Rausch, L.D. Riddick, C. Schoch, A. Shkeda, S.S. Storz, H. Sun, F. Thibaud-Nissen, I. Tolstoy, R.E. Tully, A.R. Vatsan, C. Wallin, D. Webb, W. Wu, M.J. Landrum, A. Kimchi, T. Tatusova, M. DiCuccio, P. Kitts, T.D. Murphy, and K.D. Pruitt. 2016. Reference sequence (RefSeq) database at NCBI: current status, taxonomic expansion, and functional annotation. *Nucleic Acids Res.* 44:D733–D745. doi:10.1093/nar/gkv1189.

Perkins, D.N., D.J. Pappin, D.M. Creasy, and J.S. Cottrell. 1999. Probability-based protein identification by searching sequence databases using mass spectrometry data. *Electrophoresis*. 20:3551–3567. doi:10.1002/(SICI)1522-2683(19991201)20:18<3551::AID-ELPS3551>3.0.CO;2-2.

Reese, M.G., F.H. Eeckman, D. Kulp, and D. Haussler. 1997. Improved Splice Site Detection in Genie. *J. Comput. Biol.* 4:311–323. doi:10.1089/cmb.1997.4.311.

Ruzicka, L., D.G. Howe, S. Ramachandran, S. Toro, C.E. Van Slyke, Y.M. Bradford, A. Eagle, D. Fashena, K. Frazer, P. Kalita, P. Mani, R. Martin, S.T. Moxon, H. Paddock, C. Pich, K. Schaper, X. Shao, A. Singer, and M. Westerfield. 2018. The Zebrafish Information Network: new support for non-coding genes, richer Gene Ontology annotations and the Alliance of Genome Resources. *Nucleic Acids Res.* 47:D867–D873. doi:10.1093/nar/gky1090.

Scott, A.F., C.A. Bocchini, J.S. Amberger, and A. Hamosh. 2018. OMIM.org: leveraging knowledge across phenotype–gene relationships. *Nucleic Acids Res.* 47:D1038–D1043. doi:10.1093/nar/gky1151.

Shapiro, M.B., and P. Senapathy. 1987. RNA splice junctions of different classes of eukaryotes: sequence statistics and functional implications in gene expression. *Nucleic Acids Res.* 15:7155–7174.

The UniProt Consortium. 2016. UniProt: the universal protein knowledgebase. *Nucleic Acids Res.* 45:D158–D169. doi:10.1093/nar/gkw1099.

Thorvaldsdóttir, H., J.T. Robinson, and J.P. Mesirov. 2013. Integrative Genomics Viewer (IGV): high-performance genomics data visualization and exploration. *Brief. Bioinform.* 14:178–192. doi:10.1093/bib/bbs017.

Wang, Y., X. Zhang, H. Zhang, Y. Lu, H. Huang, X. Dong, J. Chen, J. Dong, X. Yang, H. Hang, and T. Jiang. 2012. Coiled-coil networking shapes cell molecular machinery. *Mol. Biol. Cell*. 23:3911–3922. doi:10.1091/mbc.e12-05-0396.

Yeo, G., and C.B. Burge. 2004. Maximum entropy modeling of short sequence motifs with applications to RNA splicing signals. *J. Comput. Biol. J. Comput. Mol. Cell Biol.* 11:377–394. doi:10.1089/1066527041410418.

Zerbino, D.R., P. Achuthan, W. Akanni, M.R. Amode, D. Barrell, J. Bhai, K. Billis, C. Cummins, A. Gall, C.G. Girón, L. Gil, L. Gordon, L. Haggerty, E. Haskell, T. Hourlier, O.G. Izuogu, S.H. Janacek, T. Juettemann, J.K. To, M.R. Laird, I. Lavidas, Z. Liu, J.E. Loveland, T. Maurel, W. McLaren, B. Moore, J. Mudge, D.N. Murphy, V. Newman, M. Nuhn, D. Ogeh, C.K. Ong, A. Parker, M. Patricio, H.S. Riat, H. Schuilenburg, D. Sheppard, H. Sparrow, K. Taylor, A. Thormann, A. Vullo, B. Walts, A. Zadissa, A. Frankish, S.E. Hunt, M. Kostadima, N. Langridge, F.J. Martin, M. Muffato, E. Perry, M. Ruffier, D.M. Staines, S.J. Trevanion, B.L. Aken, F. Cunningham, A. Yates, and P. Flicek. 2017. Ensembl 2018. *Nucleic Acids Res.* 46:D754–D761. doi:10.1093/nar/gkx1098.
